# Genome-wide associations of fitness components reveal antagonistic pleiotropy and sexual conflict in the Florida Scrub-Jay

**DOI:** 10.1101/2025.09.09.673786

**Authors:** Elissa J. Cosgrove, Angela Tringali, John W. Fitzpatrick, Reed Bowman, Andrew G. Clark, Nancy Chen

**Author notes:** co-senior authors (Andrew G. Clark, Nancy Chen).

## Abstract

Natural populations often exhibit substantial genetic variation in fitness-related traits, providing the raw material for adaptation. The persistence of high levels of genetic variation in the presence of genetic drift and directional selection has long puzzled evolutionary biologists^1-3^. Recent studies suggest that antagonistic pleiotropy may play a greater role than previously appreciated, highlighting the need for more empirical studies that consider both sexual antagonism and life-history trade-offs^4^. Here, we performed a comprehensive assessment of fitness trade-offs within and between sexes in an extensively monitored natural population of Florida Scrub-Jays (*Aphelocoma coerulescens*)^5^. We found significant additive genetic variance for female fitness and identified several loci associated with variation in survival, fecundity, and lifetime reproductive success. Many loci associated with multiple fitness components had antagonistic effects, particularly within males. Considering genome-wide polygenic effects, we found multiple genetic trade-offs both within and between sexes, especially when considering different fitness components between sexes. Comparing allele frequency trajectories for putatively pleiotropic loci demonstrated that antagonistic SNPs had smaller allele frequency changes than concordant SNPs. Our fine-scale dissection of fitness components reveals substantial antagonistic pleiotropy that affects short-term evolution, highlighting the importance of antagonistic selection in maintaining genetic variation over short time scales in nature.

## Main

A central goal of evolutionary biology is to understand the causes of variation in individual fitness in natural populations^3,6^. Fitness, an individual’s expected contribution to the next generation, is a composite trait that can be partitioned into multiple functionally distinct components, including survival, breeding success, and fecundity^7,8^. Genetic variation in fitness and its components forms the raw material for adaptation by natural selection, and natural populations maintain surprisingly high levels of genetic variation in fitness^9^. Despite extensive theoretical and empirical work exploring potential explanations for this phenomenon, the relative importance of different factors that maintain genetic variation in fitness in nature remains a longstanding question in evolutionary biology^1,2^.

One potential mechanism for maintaining variation is antagonistic pleiotropy, or genetic trade-offs, which arise when alleles enhance one fitness component at the detriment of another^2,10^. Genetic trade-offs can occur between different life stages such as survival and reproduction^11,12^. These trade-offs play a key role in the evolution of life-history traits^7^ and of senescence^13^. While there are now multiple empirical examples of antagonistic selection or antagonistic genetic variation in diverse plant and animal species (e.g., ^14-22^), empirical identification of pleiotropic loci in natural populations remains challenging.

Classical population genetics theory largely dismissed antagonistic pleiotropy as a likely factor maintaining genetic variation in natural populations, except under fairly restrictive scenarios^11,23,24^. However, recent theoretical work suggests trade-offs have greater potential for contributing to the maintenance of variation, emphasizing the need for more empirical studies that test for different forms of trade-offs^4,7,10,14,22,25,26^. Accurately measuring multiple fitness components for large numbers of individuals in natural populations is challenging, and few empirical studies test for antagonistic pleiotropy across both life stages and sexes (but see Wong & Holman^27^). To our knowledge, no study to date has performed a detailed assessment of all components of fitness and potential forms of genetic trade-offs in a wild population.

Here, we perform a comprehensive, genome-wide assessment of genetic trade-offs between different fitness components in the Florida Scrub-Jay (*Aphelocoma coerulescens*). A population of individually-marked Florida Scrub-Jays at Archbold Biological Station has been studied for decades, resulting in detailed life histories and lifetime fitness measures for thousands of individuals on a multigenerational pedigree^5^. The extensive monitoring of this natural population allows us to quantify variation in both lifetime reproductive success (LRS), defined here as the total number of offspring produced by an individual, as well as several components of fitness. We considered eleven fitness measures in males and females separately, including multiple measures of survival early and late in life, breeding success, and fecundity (Figure 1). We estimated levels of additive genetic variance for each fitness measure and performed genome-wide associations to identify loci associated with variation in fitness and its components. We systematically tested for different genetic trade-offs genome-wide and assessed the contributions of different fitness components to variation in total fitness. Finally, we compared patterns of observed allele frequency change for candidate concordant and antagonistic pleiotropic loci to explore the effects of antagonism on short-term evolutionary change.

**Figure 1.**
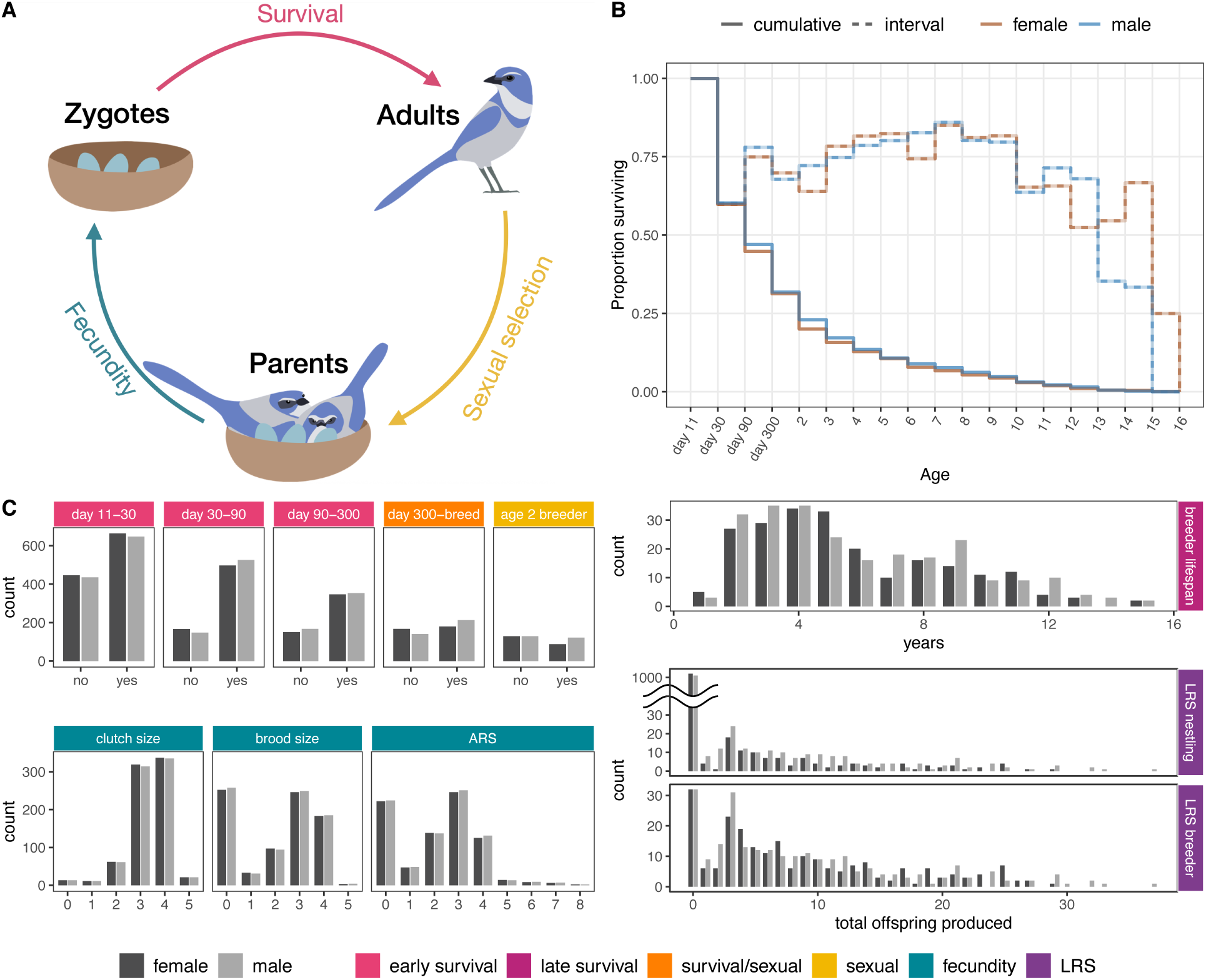
Variation in fitness in the Florida Scrub-Jay. **(A)** The life cycle of the Florida Scrub-Jay and fitness components. Illustrations by Jeremy Summers. **(B)** Cumulative survival (solid lines) of female (brown) and male (blue) Florida Scrub-Jays and the probability of survival within each interval (dashed lines). Note that the first four intervals correspond to important developmental milestones in days, and the remaining intervals correspond to age in years. **(C)** Sex-specific observation counts for all considered fitness measures. For survival and sexual selection measures, we show the number of individuals who did or did not survive a given time period or establish as a breeder, respectively. ARS: annual reproductive success, measured as the count of offspring who survived to 11 days of age produced in a year; LRS: lifetime reproductive success, measured as the count of day 11 offspring produced over a lifetime. For visual clarity, the LRS_nestling_ plot includes a jump in the *y*-axis for the bars at LRS_nestling_ = 0 (female = 1,040; male = 1,033). See Extended Data Table 1 for descriptions of each fitness measure.

### Measuring fitness and its components in the wild

From extensive individual-based field data, we quantified nine measures of fitness components and two measures of total fitness (Figure 1). We considered four measures of survival across the lifespan of a Florida Scrub-Jay. We measured early survival across three non-overlapping intervals between important milestones in juvenile development: day 11 post-hatching (when individuals are first banded and therefore can be tracked) to day 30 (when individuals are capable of sustained flight), day 30 to day 90 (when individuals reach nutritional independence), and day 90 to day 300 (when individuals are physiologically capable of breeding)^28^. We measured late survival as the lifespan of individuals who bred. Survival rates did not differ between females and males for any of the considered measures (Fisher’s Exact Test *P* > 0.05 for each early survival measure; Kolmogorov-Smirnov test *P* = 0.65 for late survival). To assess the potential for sexual selection, we considered two measures of breeding success: whether an individual was a breeder at age 2 and whether individuals who survived to day 300 established as a breeder (a blended measure of sexual selection and survival). Males had higher success rates for the latter measure (Fisher’s Exact Test *P* = 0.033), likely partially due to female-biased dispersal^29^. Finally, we considered three measures of fecundity: clutch size (number of eggs laid) and brood size (number of hatched eggs) for the first clutch of a given breeding season, and annual reproductive success (ARS), measured as the number of offspring surviving to day 11 in a year. The vast majority of first clutches had 3 or 4 eggs, while there was greater variation in brood size and ARS (Figure 1C).

In addition to detailed analysis of fitness components across the life cycle, we also quantified variation in total fitness (LRS, measured as the total day 11 offspring produced in an individual’s lifetime) in two groups of individuals: LRS_nestling_ included all individuals born in our study area who survived to 11 days of age (and therefore could be individually tracked throughout their lifetimes), and LRS_breeder_ included all breeding individuals, regardless of where they were born (and thus included immigrant breeders). LRS varied widely in our population, with a maximum LRS of 37 (Figure 1C). Most individuals (88%) never produced any offspring, which is unsurprising given only 31% of day 11 nestlings survived to adulthood (age 1 year) and 18% successfully became breeders. As expected, variation in LRS_breeder_ is less zero-inflated: only 32 of 220 (15%) female and 32 of 240 (13%) male breeders never produced any offspring. LRS_nestling_ values were lower for females than males (Kolmogorov-Smirnov test: LRS_nestling_ *P* = 0.040; LRS_breeder_ *P* = 0.779), again likely due to female-biased dispersal. As LRS is a composite trait composed of different fitness components, we were not surprised to find multiple positive phenotypic correlations between LRS and individual fitness components (Supplemental Figure 1). LRS_nestling_ is strongly correlated with day300-breed and breeder lifespan, and LRS_breeder_ is most correlated with breeder lifespan, suggesting that the primary driver of variation in total fitness in our study population is whether an individual successfully obtains breeder status and then its lifespan as a breeder.

### Determinants of fitness variation in a natural population

Individual fitness and fitness components are complex traits that are affected by a myriad of genetic and environmental factors. Given extensive knowledge of the natural history of Florida Scrub-Jays and detailed data on spatial and temporal variation in habitat, climate, and social environment (Extended Data Figure 1, Supplemental Figure 2), we were able to control for environmental heterogeneity in our analyses. We did not consider individual morphometric measures and incubation date because of possible genetic contributions to these variables and overlaps with other environmental covariates (Supplemental Text; Supplemental Figures 3-4). We used a subsampling approach^30^ to identify factors that contributed to variation in each of our sex-specific fitness measures, which we then included in all subsequent models. Consistent with previous work this step identified associations between multiple fitness measures and several measured factors, including inbreeding, territory size, and helper count^5,28,31-33^ (Supplemental Text).

With known covariates of fitness in hand, we then used Bayesian generalized linear mixed models known as “animal models”^34,35^ to partition variation in each fitness measure into different components and to test for significant additive genetic variance (Figure 2). The proportion of phenotypic variance due to additive genetic variance represents the narrow-sense heritability of the trait, which is an important predictor of the population’s potential to respond to selection^36^. We found evidence for an additive genetic variance contribution in female day30-90 survival and female LRS. Variation in early life (day11-30) survival for both females and males was largely driven by the natal nest environment, consistent with Florida Scrub-Jay fledging times and the often shared fate of offspring in the early stages of life. We observed significant cohort effects for multiple fitness components, indicating that temporal environmental variation plays an important role in variation in survival and reproduction. The identity of the female mate contributed significantly to variation in male fecundity, as we might expect given the physiological contribution of females to reproduction. Finally, fixed effects contributed significantly to variation in male and female early survival, breeder status, clutch size, and ARS, explaining as much as 18-23% of phenotypic variance in whether age 2 individuals are breeders (Extended Data Figure 2). Given the complexity of mate choice and territory acquisition in Florida Scrub-Jays^5^, it is unsurprising that environmental covariates have large contributions to sexual selection outcomes in this species.

**Figure 2.**
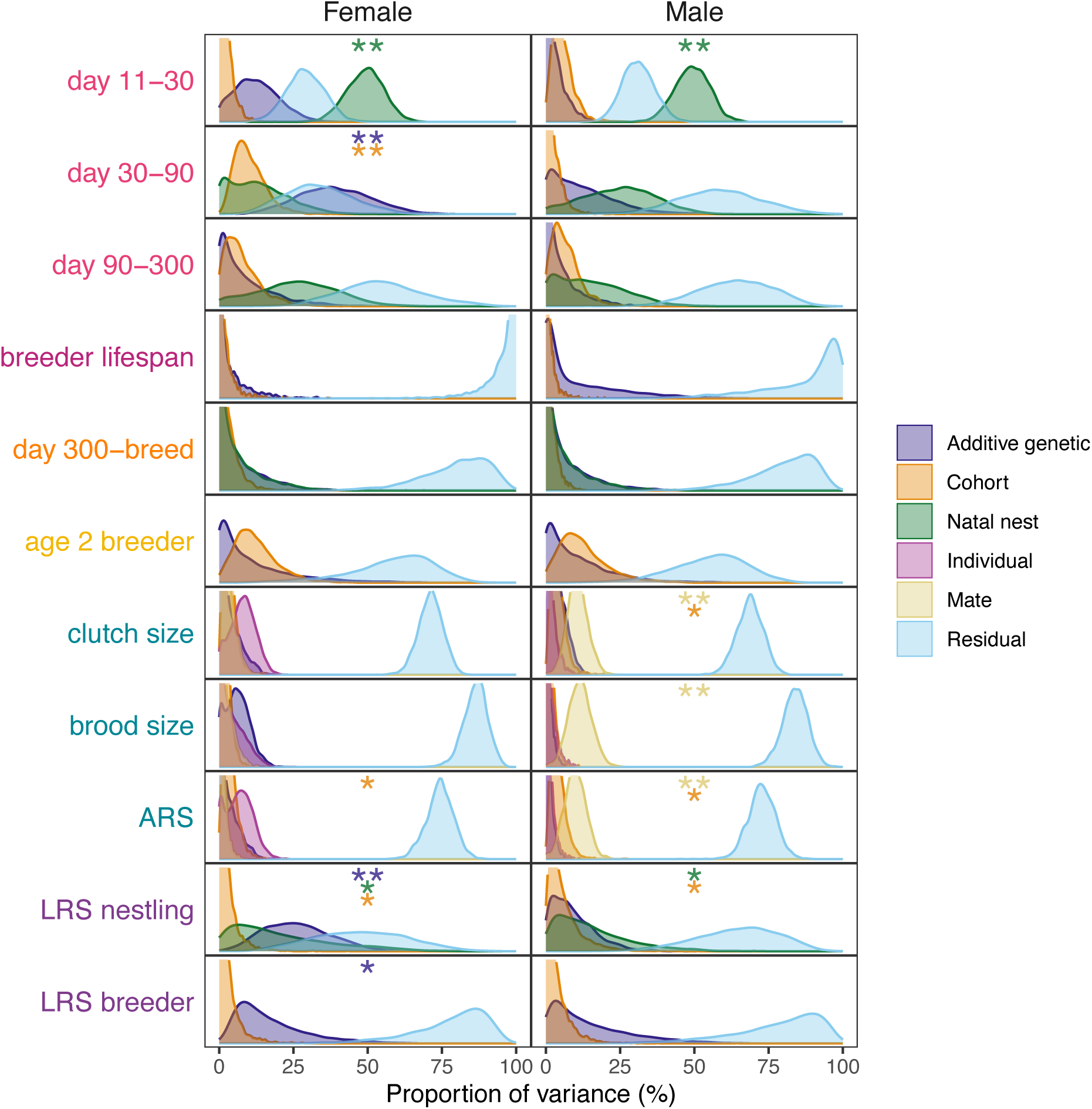
Components of variance in female and male fitness components and lifetime reproductive success. Posterior distributions of additive genetic variance (purple) as well as variance due to cohort (orange) or natal nest (green) effects. Cohort effects refer to breeding cohort for fecundity measures and natal cohort for all other analyses. For fecundity analyses, individual ID and mate ID are shown in pink and yellow, respectively. Single and double asterisks indicate that the lower bound of the 95% credible interval of corresponding variance estimate is greater than 0.001 or 0.01, respectively.

### Mapping loci associated with fitness

To estimate the effect of each SNP on each fitness measure, we performed a series of genome-wide association analyses with both additive and dominance genotype encodings. We identified 16 SNPs that were significantly associated with a fitness component and 6 SNPs significantly associated with LRS with a per-analysis Benjamini–Hochberg adjusted *P* < 0.1 (Extended Data Tables 2 & 3; Extended Data Figures 3 & 4). None of the LRS-associated SNPs were in linkage disequilibrium (LD) with SNPs associated with specific fitness components. We found the highest number of significant associations for female fecundity (12 hits across three female fecundity measures). Overall, SNPs with relatively large effects on fitness are distributed across the genome (Extended Data Figures 3 & 4). We identified 13 SNPs in or near genes (1 exonic, 6 intronic, 1 UTR, 5 within 5 kbp up- or downstream), including all 6 of the LRS-associated SNPs (Extended Data Tables 2 & 3). The small number of large effect loci associated with any fitness measure is consistent with our expectation that fitness and its components are highly polygenic traits.

### Identification of putatively pleiotropic SNPs

We next identified putatively pleiotropic loci and categorized them as antagonistic (effect size estimates in opposite directions) or concordant (effect size estimates in the same direction). This analysis excluded measures of total fitness and excluded within-sex comparisons of measures of the same fitness component. As our genome-wide association analyses are unable to detect all true associated SNPs, we used a broader dataset here to better capture overall patterns of pleiotropy. Using a nominal significance threshold of *P* < 0.01, 12.9% of all LD-pruned SNPs that were associated with at least one fitness component (2,805) had associations with two or more different fitness components within a given sex or in one or more fitness measures in both sexes (Figure 3). We found more antagonistic than concordant SNPs in males (binomial test *P* = 0.013), particularly for early survival vs. fecundity in males (binomial test *P* = 0.017). Using a more stringent significance threshold of *P* < 0.001, we found three concordant SNPs and six antagonistic SNPs (Extended Data Figure 5), most of which were observed in comparisons of late survival and fecundity (2/3 concordant SNPs and 3/6 antagonistic SNPs). A more detailed breakdown of antagonistic and concordant pleiotropic SNPs across our nine fitness components and two total fitness measures reveals potential sex differences in life history trade-offs that may reflect differences in reproductive effort: SNPs associated with clutch size in both males and females mostly have antagonistic effects on female breeder lifespan and both measures of LRS but concordant effects on male breeder lifespan (Extended Data Figure 6). Our results suggest there is substantial antagonistic pleiotropy between life stages, especially between early survival and fecundity within males.

**Figure 3.**
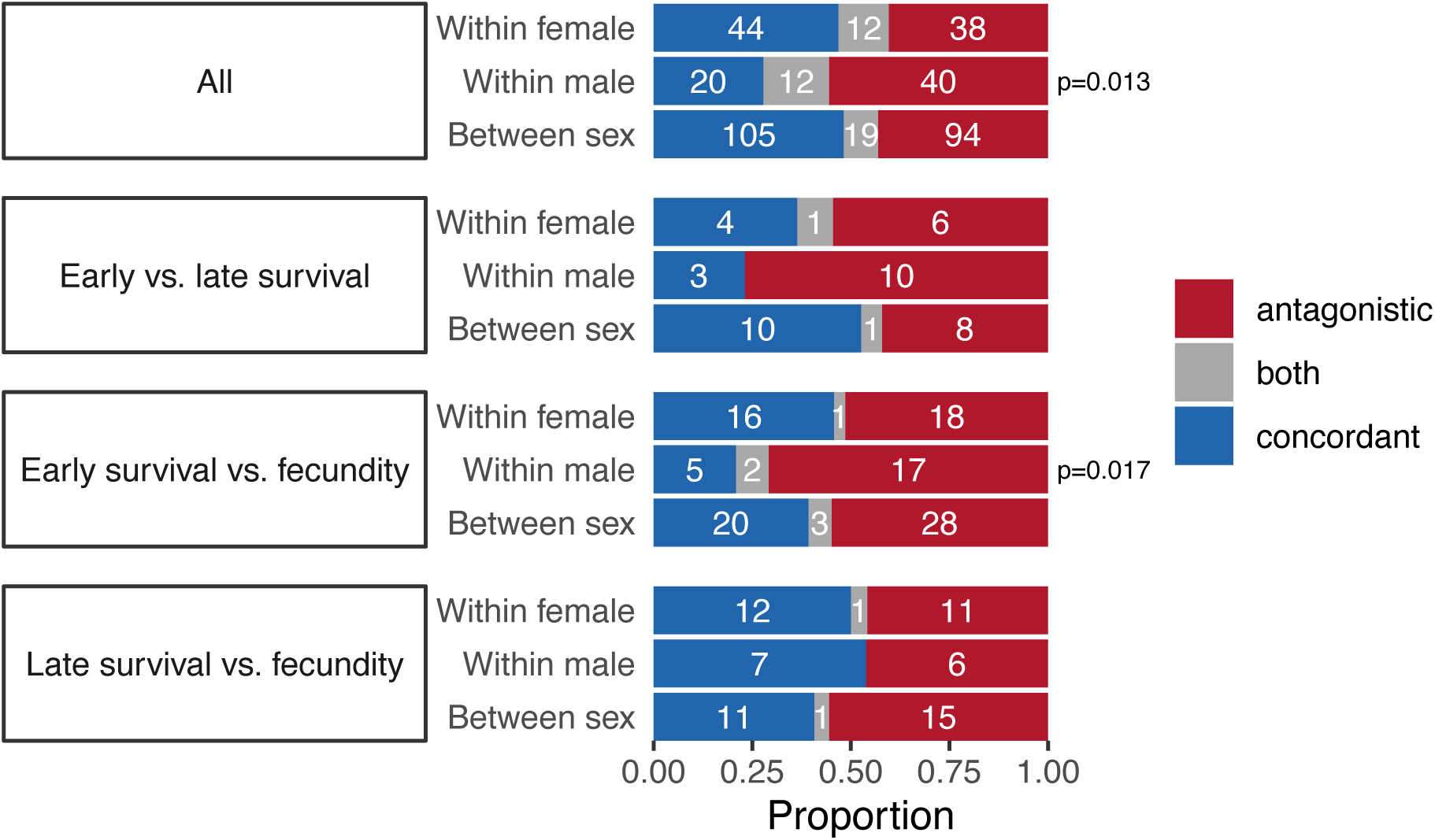
Counts of putatively pleiotropic SNPs, stratified by sex and fitness component. Counts of antagonistic (red) and concordant (blue) pleiotropic SNPs within or between sexes across all fitness components or for specific fitness component comparisons. SNPs that had both antagonistic and concordant effects are indicated in gray. A binomial test of antagonistic vs. concordant SNP counts was applied to each row (excluding SNPs with both effects) and tests with *P* < 0.05 are shown along the right side of the plot.

### Genome-wide polygenic signatures of antagonistic pleiotropy

As we expect fitness and its components to be highly polygenic traits, we also explored correlations of genome-wide *z*-scores of effect sizes for all LD-pruned SNPs for different fitness measures within and between sexes. We report significant correlations with a Benjamini-Hochberg adjusted *P* < 0.1 (Figure 4). Overall, we found more antagonistic pleiotropy in males compared to females: while 8 of 40 significant correlations between fitness measures are negative in females, 16 of 40 correlations are negative in males.

**Figure 4.**
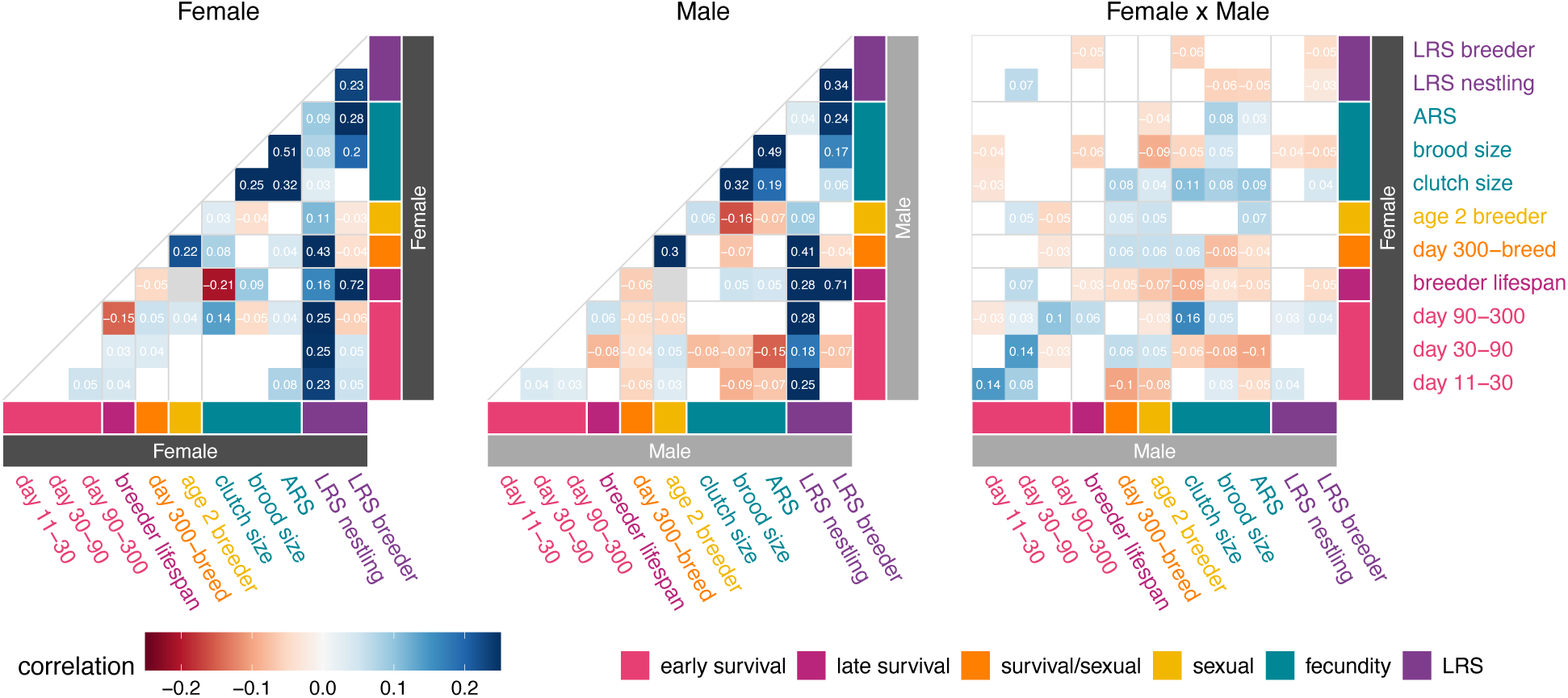
Polygenic signatures of pleiotropy show substantial genome-wide antagonism. Pairwise Spearman correlation between effect size *z*-scores for 7,998 LD-pruned SNPs for each pair of fitness measures within females (left), within males (center), or between sexes (right). Correlation *P*-values were generated via block jackknife (100 blocks). Only correlations with Benjamini-Hochberg adjusted *P* < 0.1 are shown. The correlation color scale ranges from -0.25 to 0.25 (note that negative, antagonistic correlations are red).

We found evidence of genome-wide trade-offs between early- and late-survival in both sexes: breeder survival is negatively correlated with survival from day 90 to day 300 in females and survival from day 30 to day 90 in males. We found strong support for the well-established life-history trade-off between survival and fecundity both within and between sexes. In males, all significant correlations between fecundity and early survival measures are negative, and we also see some negative correlations between fecundity and survival in females. Genome-wide *z*-scores of SNP effect sizes for breeder lifespan and clutch size are negatively correlated in females, yet *z*-scores for breeder lifespan and brood size (number of hatchlings) are positively correlated in both sexes, suggesting that sex differences in parental investment can result in different life-history trade-offs in males and females^37^. In fact, higher clutch size is associated with lower survival to the next breeding season in females (β = -0.25, *P* = 0.02) but not males (β = 0.13, *P* = 0.18; see Supplemental Text). For cross-sex comparisons (Figure 4, right plot), just over half of the significant *z*-score correlations between female and male fitness measures are negative (34 of 66, or 52%). Grouping measures by fitness component, we find more sexual antagonism between different fitness components than within the same component (mean correlation between components = -0.009 and within components = 0.045; *t* = -4.10; *P* = 0.0002). Genome-wide *z*-scores for the same fitness measure between sexes tend to be positively correlated, except for breeder lifespan and LRS_breeder_. We also found evidence of sexual conflict in some measures of early life survival as well as between male clutch size and female brood size. We found small but significant negative correlations between both female LRS measures and all male fecundity measures, as well as between both male LRS measures and female brood size. We observed a negative correlation between breeder lifespan of one sex and LRS_breeder_ of the other sex, as well as between male LRS_breeder_ and both female LRS measures. Additional analyses confirmed that observed sex differences are not driven by sex-biased immigration or differences in selected fixed effects (Supplemental Text, Supplemental Figures 5-6). Analyses using only LD-pruned SNPs with *P* < 0.01 for at least one of the considered fitness measures recovered antagonism between survival and fecundity both within and between sexes (Supplemental Text, Supplemental Figure 7). Overall, our results indicate the presence of substantial life history trade-offs genome-wide, particularly within males or between sexes. Our findings also indicate sexual antagonism for overall fitness in our population, despite an absence of sexual conflict for most individual fitness components.

### Implications for maintenance of genetic variation

To assess associations between pleiotropy and allele frequency change, we calculated the frequency of the minor allele in each birth cohort we genotyped (1989-1991, 1995, 1999-2013). We fitted a linear model of arcsine square root transformed allele frequencies over time and used the estimated slope as a measure of the degree of allele frequency change during our study period. We observed significant positive correlations between effect size and allele frequency change for most but not all fitness measures (Extended Data Figure 7). Of the 363 putatively pleiotropic SNPs, antagonistic SNPs had smaller allele frequency changes than concordant SNPs (one-tailed Student’s *t*-test *P* = 0.026) (Figure 5). We also observed that neutral SNPs (defined as SNPs that never had effect size *P* < 0.01 across all analyses) had smaller allele frequency changes than concordant SNPs (one-tailed Student’s *t*-test *P* = 0.021). Of the 51 concordant SNPs with a suggestive allele frequency change (*P* < 0.01), 43 (84%) showed allele frequency changes in the same direction as the effect size estimates. We note that as our allele frequency change estimates and genome-wide association analyses use overlapping datasets, these results allow us to understand the impact of pleiotropy on allele frequency change but do not represent independent replication of the effects. Though our fitness effect size estimates are not entirely independent of allele frequency changes, we found that they are not tightly correlated for all fitness measures (Extended Data Figure 7), consistent with heterogeneity in the impact of selection across life stages and sexes. Further consideration of the separation of allele frequency change calculations from the inference of pleiotropy is given in the discussion.

**Figure 5.**
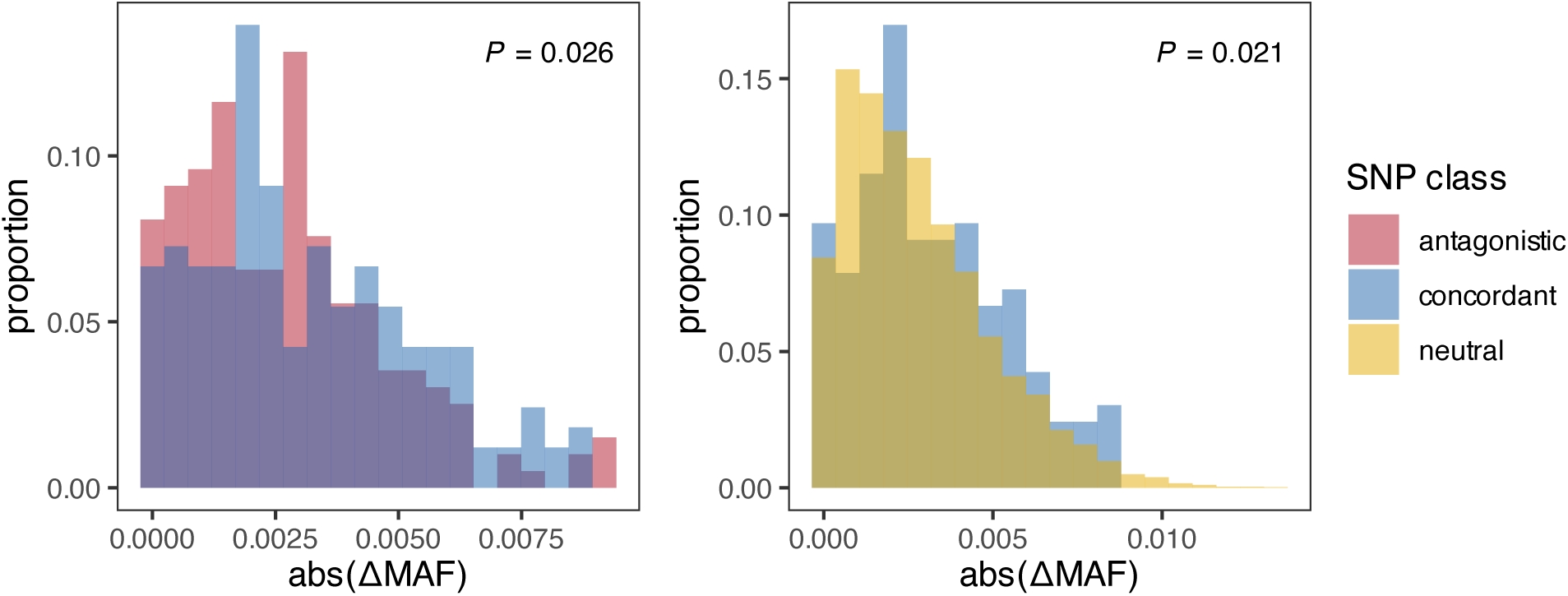
Effects of pleiotropy on short-term allele frequency change. Distributions of allele frequency change (ΔMAF) between 1989-2013 for putatively concordant (blue, *n* = 165) vs. antagonistic (red, *n* = 198) SNPs (left) and concordant vs. neutral (yellow, *n* = 7,261) SNPs (right) have significantly different means (one-tailed Student’s *t*-test *P* = 0.026 and 0.021, respectively). Histograms are semi-transparent and overlaid.

## Discussion

One of the deepest unresolved paradoxes in evolutionary biology is the maintenance of high levels of genetic variation in fitness and its components under stabilizing or directional selection in natural populations^1-3^. Here, we have conducted a comprehensive analysis of fitness and its many components in a natural population and quantified genetic trade-offs among fitness components within and between sexes. Our findings reveal additive genetic variation in fitness that differed between sexes, with a few instances of single SNPs showing association with fitness components. Analyses of both putatively pleiotropic sites and genome-wide correlations found evidence of substantial antagonistic pleiotropy, especially between survival and fecundity within males or between sexes, with pleiotropy affecting patterns of allele frequency change over time. Together, our results indicate that complex genetic trade-offs can be pervasive in the wild, with important implications for the maintenance of genetic variation and evolutionary constraints on adaptation in natural populations.

Our results suggest that fitness and its components have low heritability and are highly polygenic, consistent with theoretical expectations and previous empirical work^38,39^. We detected non-zero additive genetic variance in female day30-90 survival and LRS, with LRS estimates within the range found in Bonnet et al.^9^ Fitness traits are expected to have low heritability because they tend to be more complex and highly influenced by environmental variance^40^. Our results highlight the importance of accounting for environmental heterogeneity in fitness variation^35^, with fixed effects explaining 18-23% of the phenotypic variance in different fitness measures. Our genome-wide association analyses identified a limited number of SNPs across the genome that were significantly associated with fitness components. While the sparse detection of large-effect loci may also be influenced by statistical power limitations, low SNP density, and the upward bias of effect size estimates in studies with limited sample size^41-43^, these results are consistent with other GWAS studies of adaptive traits that report a polygenic basis for complex phenotypes like fitness (*e.g.*, ^44-48^). Most fitness-associated variants (15/22) were identified from models where the major allele is dominant rather than additive, reinforcing the need for better estimates of dominance effects in population genetic studies, as dominance can play a crucial role in shaping the genetic architecture of fitness traits^49^.

Our results suggest there may be sex differences in the genetic architecture of fitness. For instance, female day 30-90 survival showed significant heritability, while male survival in the same period did not, potentially due to male social dominance affecting resource access during critical developmental periods^28,50^. For total fitness, we found significant heritability of LRS in females but not males. Other studies have documented greater heritability for female fitness than male fitness in a variety of species^14-21,27^. While we observe sex differences in whether heritability of early survival and LRS is significantly different from zero, we note that the 95% credible intervals for each measure in the two sexes overlap, and we would need greater sample sizes to more rigorously test the significance of sex differences in heritabilities. Our results cannot be explained by errors in paternity assignments, as we used a GRM calculated with our genomic data and removed the few cases of known extra-pair paternity from our analyses (Supplemental Text). Instead, sex differences in heritability could be caused by greater female control of reproduction^51^ or potentially reflect stronger past selection on males^52^.

With estimates of genetic effects for multiple fitness components in hand, we were able to test for genetic trade-offs among fitness components within and between sexes. Analyses of both putatively pleiotropic SNPs and genome-wide correlations found genetic trade-offs between survival and reproduction as well as between early and late survival. Our results add to accumulating empirical evidence of antagonistic pleiotropy^14-22^ and provide a counterexample to recent reports of prevalent positive pleiotropy across fitness components^27^. Particularly striking was the pattern of more antagonistic pleiotropy within males compared to females, suggesting that males may face more stringent genetic constraints due to antagonistic pleiotropy^53^. This sex difference in the prevalence of genetic trade-offs may reflect differences in life-history strategies or the strength of sexual selection acting on males versus females^54^. In Florida Scrub-Jays, it is likely that the requirements for becoming a breeder are more stringent for males than females^55^. In addition to within-sex, inter-component trade-offs, our analysis revealed extensive sexual antagonism. We found evidence for sexual conflict in overall fitness despite concordant effects for most individual fitness components, though we found substantial antagonism between different fitness components between sexes. Our results suggest that sexual conflict may be more prevalent in natural populations than previously recognized^4^, possibly due to the greater opportunity for pleiotropic constraints to accumulate over extended lifespans or the sparsity of empirical studies that test multiple fitness components.

The observed genetic trade-offs have important implications for evolutionary trajectories and the maintenance of genetic variation in natural populations. While antagonistic pleiotropy can promote the persistence of polymorphism, and a small number of balanced polymorphisms can have a large impact on standing genetic variation in fitness^7^, it may also constrain adaptive responses to selection^56,57^. Analysis of allele frequency changes over a 25-year period in our study population revealed that antagonistic SNPs had significantly smaller allele frequency changes than concordant SNPs (*P* = 0.026), providing some direct empirical evidence that genetic trade-offs can slow evolutionary change and help maintain genetic variation in natural populations. This finding is consistent with theoretical predictions that antagonistic pleiotropy should reduce the rate of allele frequency change and increase the transit time for polymorphism by balancing positive and negative selection effects across different fitness components^58^. Although pleiotropy was estimated using the same genotyped cohorts, there are key differences between our measure of pleiotropy and our measure of allele frequency change: pleiotropy reflects patterns of sex-specific SNP–phenotype associations within cohorts, while allele frequency change quantifies temporal shifts in allele counts across cohorts, regardless of sex. Thus, the observed difference in frequency change between pleiotropic and non-pleiotropic SNPs is not a tautology. We also find that neutral SNPs (those with no significant fitness associations) had smaller allele frequency changes than SNPs showing concordant pleiotropy (*P* = 0.021), suggesting that even weak selection can detectably influence evolutionary dynamics over short time scales in natural populations. Future studies with higher resolution genomic data as well as further consideration of gene flow and variation in the strength and direction of selection are needed to confirm these trends.

Several caveats should be considered when interpreting our results. Differences in statistical power among fitness components, limited sample sizes, and low SNP density all constrain our ability to accurately estimate low heritabilities and small effect sizes^59^. Our modest sample sizes (*n* = 217-1210 depending on the fitness component) limited our power to detect variants with small effects on fitness, a problem that was particularly acute in estimates of sexual selection. Our relatively low SNP density (12,208 SNPs) meant that we relied on SNPs as markers for many causal effects in LD across a fraction of the genome, and we had little chance to identify causal variants through their inferred molecular disruption. Truly pleiotropic effects of single variants may also be confounded with distinct effects of pairs of variants in LD. Additionally, our analyses do not account for indirect genetic effects, which may lead to underestimation of total heritable variation^60,61^.

From a conservation perspective, our findings underscore the complexity of predicting population persistence in the face of environmental change. The genetic architecture of fitness, including the prevalence of genetic trade-offs, can influence both the maintenance of genetic variation and the capacity for adaptation^62^. The Florida Scrub-Jay is a federally Threatened species with historically low effective population sizes (*N_e_* ≈ 18 to 952 across local populations in Florida^63^, which typically results in random genetic drift overpowering natural selection at most loci^64^. The detection of significant genetic trade-offs in this small population suggests that antagonistic pleiotropy may help maintain genetic variation even in the face of strong genetic drift, potentially providing some resilience against environmental change. On the other hand, adaptive potential in at-risk populations may not only depend on levels of genetic diversity but could also be limited by genetic trade-offs. Our results highlight the need for more studies of the genetic basis of fitness in threatened species, particularly those with different life histories and mating systems.

In conclusion, our study demonstrates the power of combining extensive longitudinal individual-based field monitoring with genomic approaches to dissect the genetic underpinnings of fitness and its components and to detect genetic trade-offs in natural populations. The comprehensive nature of this study, examining 11 different fitness measures across both sexes and multiple life stages, together with 70 potentially relevant but confounding environmental variables, provides one of the most detailed portraits of fitness genetics in a natural population to date. Perhaps most surprisingly, we identified substantial antagonistic pleiotropy between life stages and sexes with corresponding associations with allele frequency change. These findings indicate that genetic trade-offs may be more pervasive in nature and play a more important role in short-term evolutionary dynamics than previously appreciated. Ongoing whole-genome sequencing will enable more detailed investigations of the molecular mechanisms underlying fitness trade-offs and their evolutionary consequences. Future research should aim to replicate these findings in other taxa and environments, considering how social structure and life history of a species may influence the relative importance of different types of genetic trade-offs. Such efforts are critical for advancing our understanding of evolutionary processes and for developing effective conservation strategies for threatened populations.

## Methods

### Study system and dataset

Florida Scrub-Jays are non-migratory and highly territorial cooperative breeding birds endemic to oak scrub in Florida, USA^5^ with low dispersal distances and high breeding site fidelity^29,65,66^. A population of Florida Scrub-Jays at Archbold Biological Station (Venus, FL, USA) has been intensively monitored since 1969^5^. Individuals are banded with unique color combinations and the entire study population is censused monthly, providing detailed information on individual lifespans as well as the ability to identify immigrants. Territories are mapped each year, accruing data on territory size and habitat composition for 45-88 territories each year. During the breeding season (March - May), all nests are found and monitored until fledging or failure, resulting in accurate measures of annual and lifetime reproductive success. Offspring are captured and banded at 11 days after hatching and again at 60-90 days after hatching. All work with live birds was approved by the Institutional Animal Care and Use Committees at Cornell University (IACUC 2010-0015) and Archbold Biological Station (AUP-006-R) and permitted by the US Fish and Wildlife Service (TE824723-8, TE-117769), the US Geological Survey (banding permits 07732, 23098), and the Florida Fish and Wildlife Conservation Commission (LSSC-10-00205).

Florida Scrub-Jays are socially monogamous with exceptionally low rates of extra-pair paternity (EPP; Supplemental Text)^67,68^, allowing accurate pedigree reconstruction from field observations. We verified parentage assignments and expanded the pedigree using genomic data collected from a previous study (3,984 individuals genotyped with custom Illumina iSelect BeadChips at 15,416 genome-wide SNPs)^31^. In this study, we focused on field observations from 1989-2015 and used a subset of the genomic data from Chen et al.^31^ We restricted our analyses to cohorts in which the majority (>67%) of individuals were genotyped (see below) and excluded any nests with known EPP (Supplemental Text). Our overall dataset included 12,208 SNPs in 2,874 individuals. Across the full dataset, LD decays to R^2^ = 0.1 at 29 kb. For some analyses, we pruned SNPs for linkage disequilibrium using ‘--indep-pairwise 50 5 0.2’ in PLINK^69^ and physical positions estimated from an earlier version of the Florida Scrub-Jay genome^70^, resulting in 7,998 SNPs. We used the most current Florida Scrub-Jay annotation^71^ and SnpEff v4.3^72^ to annotate our variants.

### Measuring fitness

The extensive field monitoring of our study population allowed us to measure lifetime reproductive success as well as multiple fitness components for females and males with high accuracy (Figure 1, Extended Data Table S1). For survival, we considered three measures of juvenile survival as well as breeder survival. Florida Scrub-Jays typically fledge or leave the nest around 18 days after hatching. Fledglings become physiologically capable of sustained flight by day 30 posthatching, nutritional independence by day 90 posthatching, and breeding by day 300 posthatching^28^. We measured juvenile survival as binary responses across three important developmental time intervals: survival from day 11 (when nestlings are first banded) to day 30 posthatching, day 30 to day 90, and day 90 to day 300. For juvenile survival, we only included nestlings born in mapped territories in years when >67% of the natal cohort was genotyped: 1989-1991, 1995, 1999-2013 (“genotyped natal cohorts”).

Individuals who attain social dominance in a territory and successfully produce at least one egg are considered breeders for the rest of their lifetimes. We used breeder lifespan (in years) as a measure of late survival. We only included individuals who were observed during each breeding season of their lifespan and were from genotyped natal cohorts. Immigrant breeders are assumed to be 2 years old during their first observed breeding season. We note that 26 individuals (5.7% of the total number of sampled breeders) were still alive in September 2022, but we retained them as their lifespans were already in the top quartile (>8 years), limiting the effect of right censoring.

Measuring sexual selection in our data set was challenging, as it is confounded with survival due to the age-dependent structure of the Florida Scrub-Jay life cycle. Here, we use breeder status to test for sexual selection. Florida Scrub-Jays can attain breeder status at varying ages, with the majority of individuals becoming breeders by age 4 (if they survive that long; 80% and 93% of 3- and 4-year-old jays in our dataset are breeders, respectively). We investigated sexual selection using two binary metrics: whether individuals who survived to day 300 posthatching became a breeder (day300-breed) and whether or not an individual was a breeder at age 2 (age2 breeder). We limited both datasets to individuals from genotyped natal cohorts. We note that day300-breed is a blended measure of both survival and breeder status, as well as emigration (disappearances of non-breeding individuals can be due to death or dispersal outside our study tract, though from annual surveys of peripheral areas we have evidence that successful emigration off the study tract is relatively rare). For both measures, we observed a smaller proportion of successful females compared to males (Figure 1C), likely a consequence of female-biased dispersal.

We considered three measures of fecundity: clutch size, brood size, and annual reproductive success (ARS). Clutch size is the number of eggs laid in the first clutch of the season, brood size is the number of hatchlings in the first clutch of the season, and ARS is the total number of day 11 offspring produced by an individual in a given breeding season. We only included observations from mapped territories in years when >67% of breeders were genotyped (1989-1992, 2003-2015). For clutch size and ARS models, we included nests with zero eggs only if both breeders had laid an egg in previous years. We note that this criterion excludes five putatively sterile individuals who bred for 6-9 years but never produced an egg.

To enable comparison of specific fitness components to overall fitness, we also conducted analyses of overall lifetime reproductive success (LRS), measured here as the total number of day 11 offspring produced by an individual over their lifetime. We considered LRS for two groups of individuals: all locally-born individuals who survived to 11 days of age (LRS_nestling_, *n* = 2,358) and the set of breeding individuals, which includes immigrant breeders (LRS_breeder_, *n* = 460). Both datasets only included individuals who were observed during each breeding season of their lifespan and were from genotyped natal cohorts.

### Consideration of environmental covariates

To more directly interrogate genetic associations, we first accounted for environmental covariates affecting fitness. Thanks to detailed, multi-decadal data collection at Archbold Biological Station, we were able to consider a wealth of environmental data, including variables describing characteristics of the cohort, habitat, individual, natal nest environment, climate, and fire history (Supplemental Text; Supplemental Table 1).

We conducted variable selection for each fitness measure separately, considering 19-53 potential covariates in each analysis. We fitted generalized linear mixed models with different error distributions (Extended Data Table 1) with the R function GMMAT::glmmkin()^73^. Random effects included the genetic relatedness matrix (GRM) as well as biologically-relevant effects such as natal or breeding cohort, natal nest, mate ID, or individual ID for repeated measures (Figure 2). We used the R function AGHMatrix::Gmatrix()^74^ to calculate the GRM from autosomal SNPs using options maf=0.05 and method=“VanRaden”. We adjusted the GRM to be positive definite using the R function corpcor::make.positive.definite()^75^. We carried out model selection using a subsampling approach^30^: we fitted models on 100 independent draws of 90% of the dataset without replacement, and computed the *P*-value of the median test statistic across these models for each potential covariate. To reduce the set of potential fixed effects, we first applied the subsampling approach to all possible single- and two-variable models. We then used *P*-values from the single-variable models to select the most significantly associated single variable for each set of related climate or fire data variables (e.g., precipitation, Southern Oscillation Index (SOI), time since fire (TSF)) or for any pair of highly correlated variables (absolute value of Pearson correlation coefficient > 0.75). Finally, we discarded all variables with *P* > 0.1 in all single- and two-variable models. Following this initial filtering step, we then applied two rounds of subsampling on models with the full set of selected covariates, selecting any covariates with *P* < 0.05 in each iteration. Variables that were selected in at least one analysis are shown in Extended Data Figure 1, and variables that were never selected are shown in Supplemental Figure 2. Our final datasets for each analysis did not include any missing data; final sample sizes can be found in Extended Data Table 1.

### Estimating additive genetic variance

Using the fixed effects selected above, we ran animal models to estimate narrow-sense heritability for each fitness measure and to assess the contributions of environmental random effects. We fitted these models with R MCMCglmm::MCMCglmm()^76^ using the SNP-derived GRM, computed as described above. We used the same error distributions for each fitness measure as before (Extended Data Table 1; we used the “threshold” family for binomial models), except for our two LRS measures. Here the flexibility of MCMCglmm enabled us to fit more appropriate zero-inflated Poisson models. We followed literature recommendations for setting priors and estimating quantitative genetic parameters (Bonnet et al.^9^ for zero-inflated Poisson models, de Villemereuil^77^ for all other models). We used Gaussian priors for all fixed effects, inverse Gamma priors for all random effects in Gaussian models, and parameter expanded priors for all random effects in threshold and zero-inflated Poisson models. We computed variance proportions relative to the total phenotypic variance, which we estimated as the sum of variance component estimates (including an estimate of the variance contributed by fixed effects). We ran all models for at least 650,000 iterations with a burn-in of 150,000 (thinning interval: 500), except the LRS_nestling_ analyses, which we ran for at least 260,000 iterations with a burn-in of 60,000 (thinning interval: 200) due to computational limitations. All models passed the stationarity test for convergence and had posterior effective sample sizes >200. We ran three replicates of each model and combined the chains of replicates for all plots and values reported herein. Estimates from all models are reported in Supplemental File 1.

### Genome-wide association analysis

We tested for associations between each SNP and each fitness measure using mixed effect models fitted with the R function GMMAT::glmmkin(). As before, all models included the GRM and other biologically-relevant random effects (Figure 2) and selected fixed effects. We considered three different genotype models: additive (*AA* = 0, *Aa* = 1, *aa* = 2), major allele dominance (*AA* = *Aa* = 0, *aa* = 1), and minor allele dominance (*AA* = 0, *Aa* = *aa* = 1). For each test, we only considered variants with a minimum of 10 observations each for at least two genotype values. We computed the Wald test *P*-value for each SNP effect size estimate and adjusted *P*-values using the Benjamini-Hochberg correction for each analysis (sex x fitness measure x genotype model). We used a significance threshold of adjusted *P* < 0.1. We confirmed that raw *P*-values for each analysis followed a (roughly) uniform distribution using quantile-quantile plots (Supplemental Figure 8). In the cross-sex comparison of correlations within and between different fitness components, we grouped fitness measures by component: three early survival, one late survival, two breeder success (age2 breeder and day300-breed), and three fecundity measures.

### Identifying putatively pleiotropic loci

We defined putatively pleiotropic SNPs as SNPs that had suggestive effects (raw *P* < 0.01) in two or more fitness components. We excluded within-sex comparisons of the same fitness component (*i.e.*, we excluded comparisons between the three early survival measures and between the three fecundity measures) as well as within-sex comparisons of age 2 breeder and day 300-breed (which are inherently correlated; Supplemental Figure 1). We considered comparisons between LRS and individual fitness components separately. For any identified putatively pleiotropic SNPs in LD (R^2^ > 0.2), we only retained the SNP with the lowest *P*-value. We then counted the number of putatively pleiotropic SNPs with effect size estimates in opposite directions (antagonistic) vs. effect size estimates in the same direction (concordant) for each pair of analyses as well as for specific fitness component or sex comparisons. SNPs that showed both antagonistic and concordant effects were either considered separately (in discussion of pleiotropy) or categorized as an antagonistic SNP (for allele frequency analyses).

### Polygenic signals of pleiotropy

To explore polygenic signals of pleiotropy, we computed Spearman correlation coefficients between effect size z-scores for all LD-pruned SNPs (*N* = 7,998) from the additive genotype model for all pairs of fitness measures (including LRS) within and between sexes. We used a block jackknife approach to compute correlation *P*-values^30^, dividing position-ordered SNPs into 100 blocks. We only report correlations with a Benjamini-Hochberg adjusted *P* < 0.1.

### Change in minor allele frequency

As we have genomic data from the majority of the birth cohorts for multiple years (1989-1991, 1995, 1999-2013), we could estimate allele frequency change in our study population over a 25-year time period. We calculated the minor allele frequency for each autosomal SNP in each genotyped natal cohort. To correct for biases in allele frequency change associated with different starting allele frequencies, we applied an arcsine square root transformation to the annual minor allele frequency estimates. We then fitted a linear model of the transformed MAF vs. time for each SNP and used the estimated slope as a measure of allele frequency change. We used one-tailed Student’s *t*-tests to test for significant differences in allele frequency changes between different groups of SNPs.

### Statistical analysis

We performed all statistical analyses in R 4.0.5 and created plots in R 4.3.2^78^. We conducted Fisher’s Exact Tests, Kolmogorov-Smirnov Tests, and Student’s *t*-tests with a two-sided alternative hypothesis unless otherwise noted.

## Acknowledgments

Thank you to the many staff, interns, and volunteers at Archbold Biological Station who collected field data over the years. Thank you to Steve Schoech and his team for sharing data from the South Tract, which helped inform our pedigree and some breeder status measures. Thank you to the Cornell BioHPC for computational help and to Andrew Siefert, James Booth, and Stephen Parry from the Cornell Statistical Consulting Unit, as well as Timothée Bonnet, Doc Edge, and Graham Coop for advice on statistical analyses. Additional thanks to Graham for providing helpful comments on the manuscript. This work was funded by NSF Grants DEB-0855879, DEB-1257628, BSR-9021902, and DEB-0316292; the Cornell Lab of Ornithology and its Athena Fund; NSF Postdoctoral Fellowship in Biology 1523665; and NIH grant 1R35GM133412.

## Author contributions

R.B. and J.W.F. supervised field data collection. E.J.C., A.T., and N.C. compiled field data for analysis. E.J.C. performed data analysis with guidance from N.C. and A.G.C. E.J.C., N.C., and A.G.C wrote the manuscript, with feedback from all authors.

## Competing interests

The authors declare no competing interests.

## Additional information

Supplementary Information is available for this paper. Correspondence and requests for materials should be addressed to Nancy Chen, Elissa Cosgrove, and Andrew Clark.

---

**Extended Data Figure 1.**
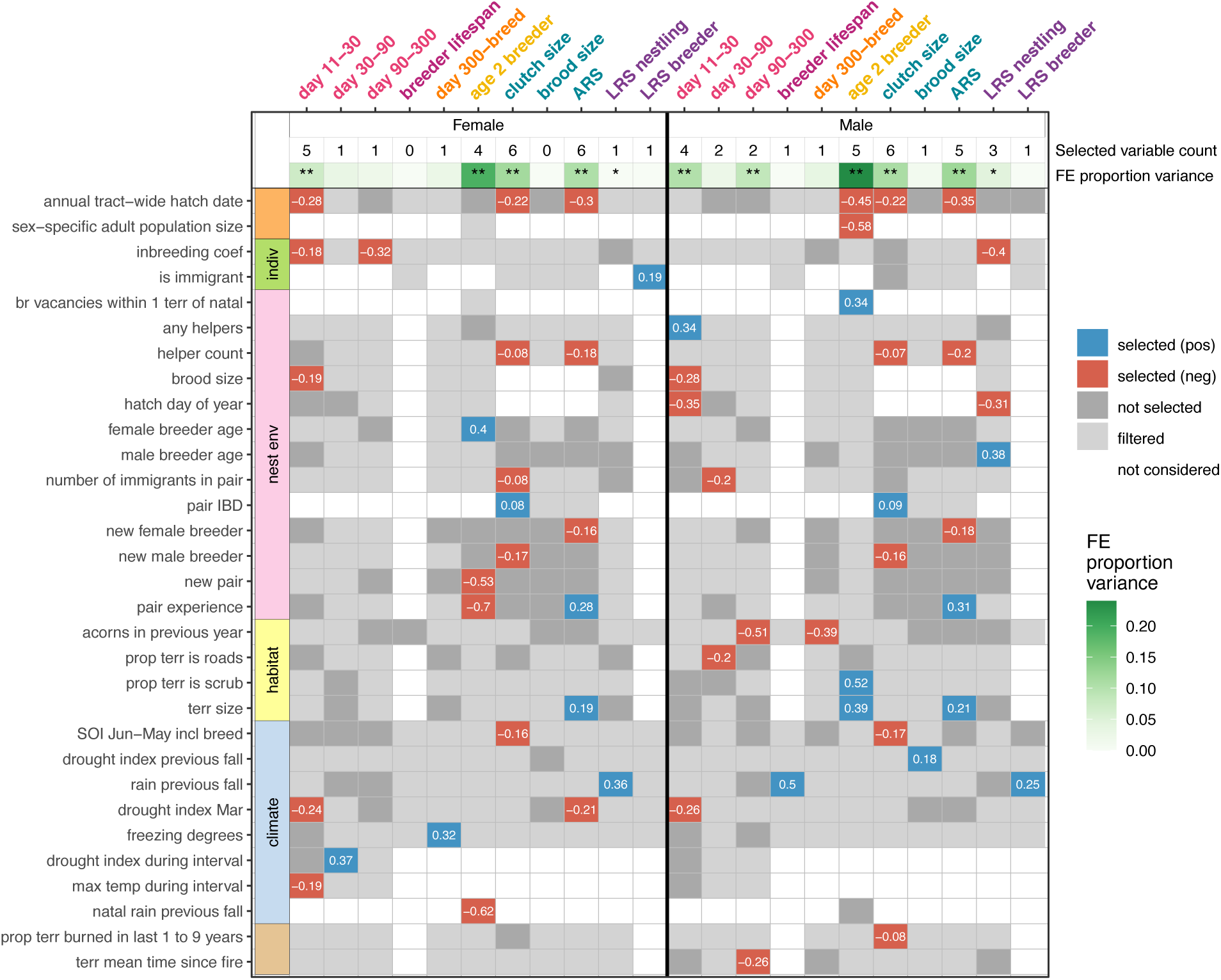
Fixed effects (FE) selected for each analysis. Numeric values indicate the regression coefficient in the final model for each fitness measure in females and males. We considered several different potential covariates (fixed effects; FE), including characteristics of the cohort (orange), individual (green), natal nest environment (pink), habitat (yellow), climate (blue), and fire history (brown). Gray shading indicates variables that were removed due to earlier filtering steps (light gray) or at the final variable selection step (dark gray). The number of variables included in downstream analyses and the posterior mode proportion of phenotypic variance explained by these variables derived from Bayesian animal models (also see Figure 2, Extended Data Figure 2) are displayed across the top. Single and double asterisks indicate that the lower bound of the 95% credible interval for the corresponding variance estimate is greater than 0.001 or 0.01, respectively.

**Extended Data Figure 2.**
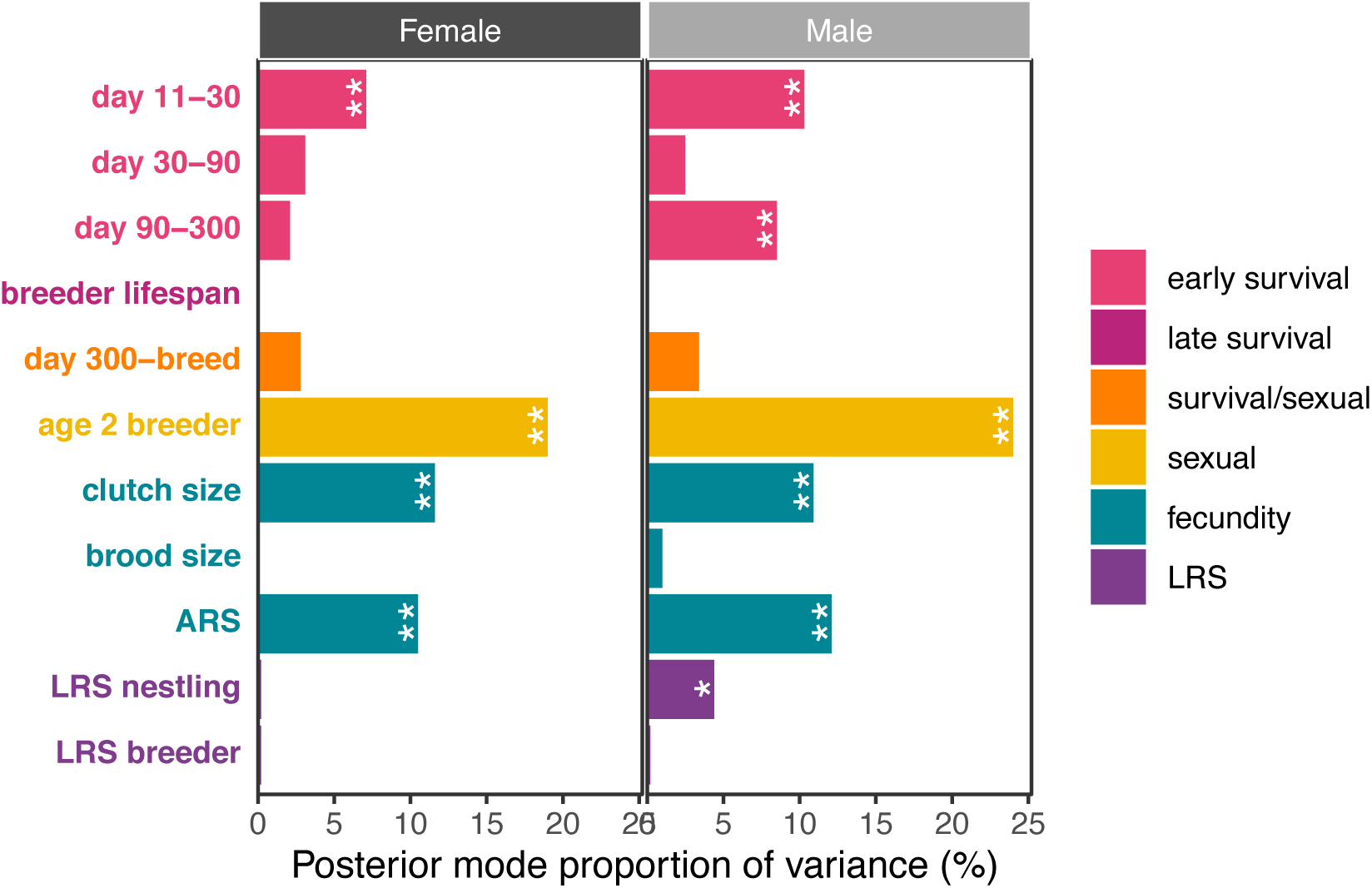
Fitness variance due to fixed effects. The phenotypic variance contributed by fixed effects across MCMC samples. Single and double asterisks indicate that the lower bound of the 95% credible interval for the corresponding variance estimate is greater than 0.001 or 0.01, respectively. See Figure 2 for other variance components estimated from these models.

**Extended Data Figure 3.**
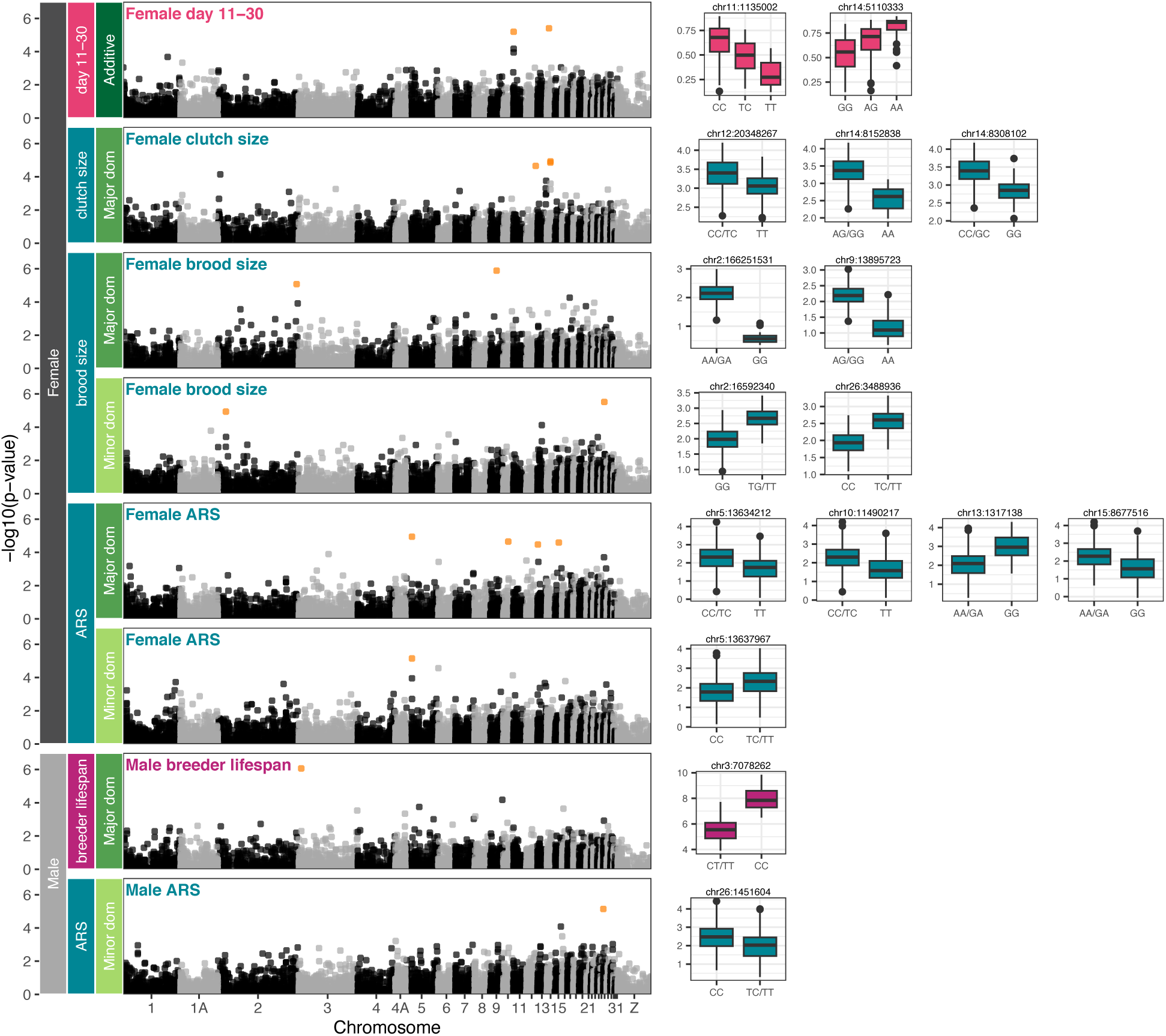
SNPs associated with fitness components. Genome-wide association analyses identified 16 SNPs significantly associated with a fitness component. Manhattan plot for each analysis with significant SNPs (Benjamini–Hochberg adjusted *P* < 0.1) in orange. Boxplots on the right show fitted values for each genotype for each significant SNP. Green labels indicate genotype model. Major dom: major allele dominance. Minor dom: minor allele dominance.

**Extended Data Figure 4.**
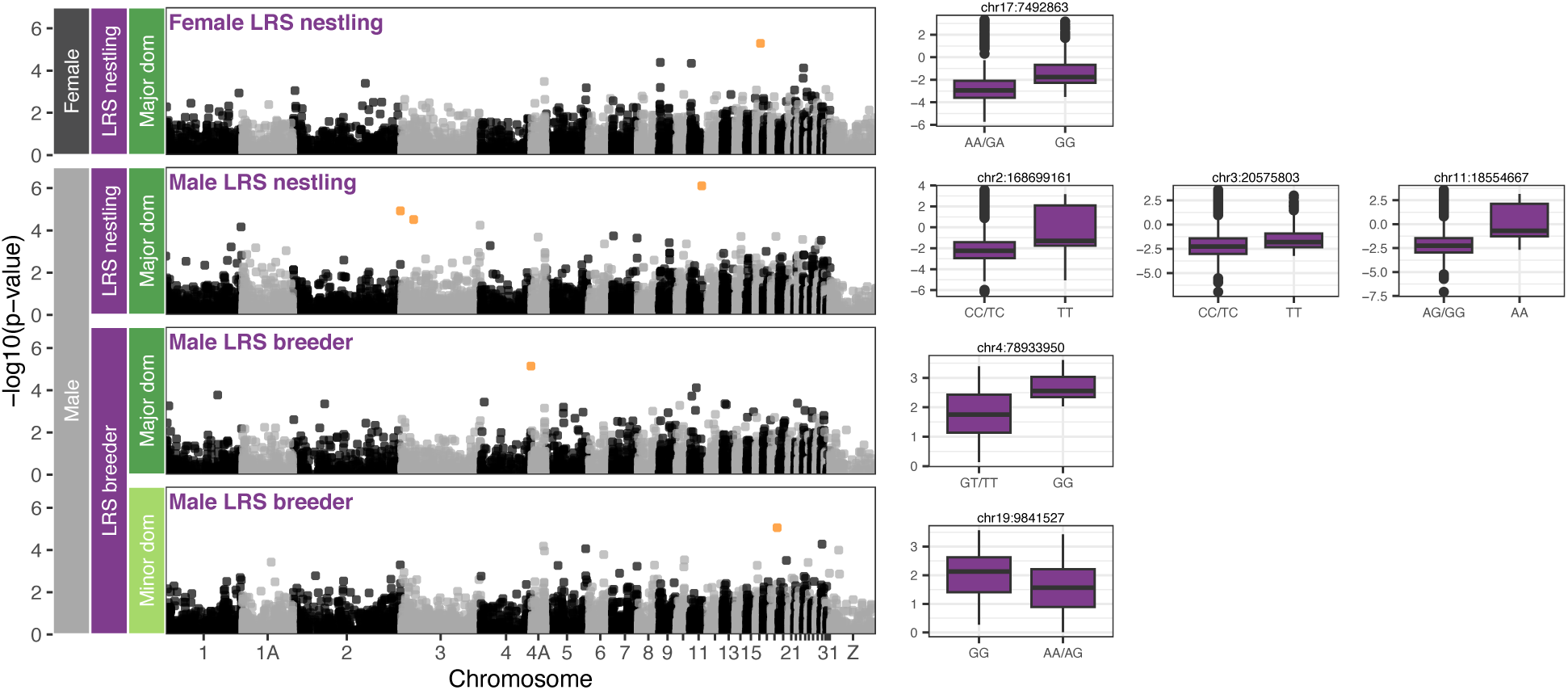
SNPs associated with lifetime reproductive success (LRS). Genome-wide association analyses identified 6 SNPs significantly associated with variation in lifetime reproductive success. Manhattan plot for each analysis with significant SNPs (Benjamini–Hochberg adjusted *P* < 0.1) in orange. Boxplots on the right show fitted values for each genotype for each significant SNP. Green labels indicate genotype model. Major dom: major allele dominance. Minor dom: minor allele dominance.

**Extended Data Figure 5.**
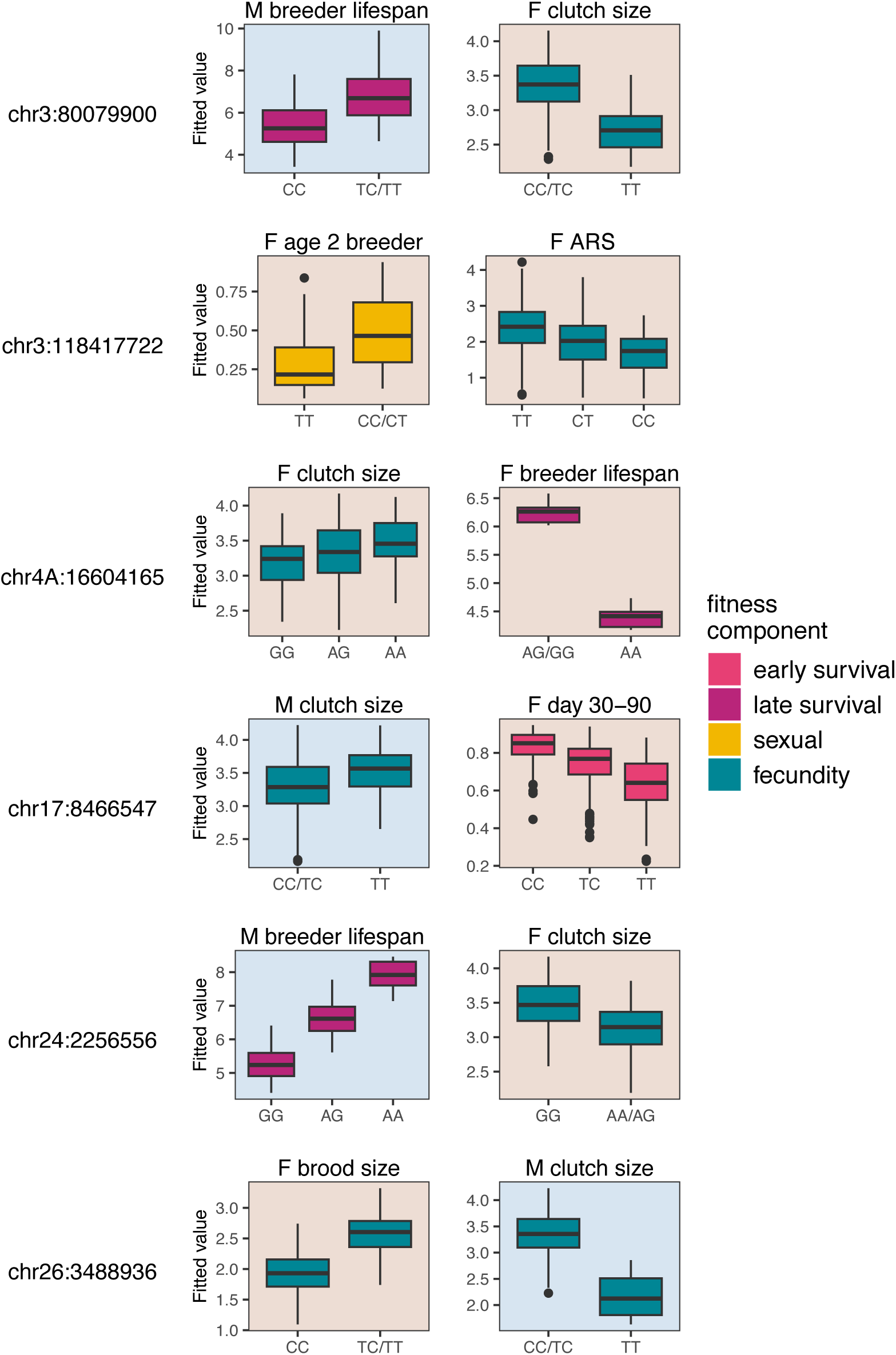
Predicted values for the strongest antagonistic SNPs. Boxplots of fitted values vs. SNP genotype for antagonistic SNPs identified using a significance threshold of *P* < 0.001. Plot background color indicates sex (Female (F): light orange; Male (M): light blue) and boxplot color indicates fitness component.

**Extended Data Figure 6.**
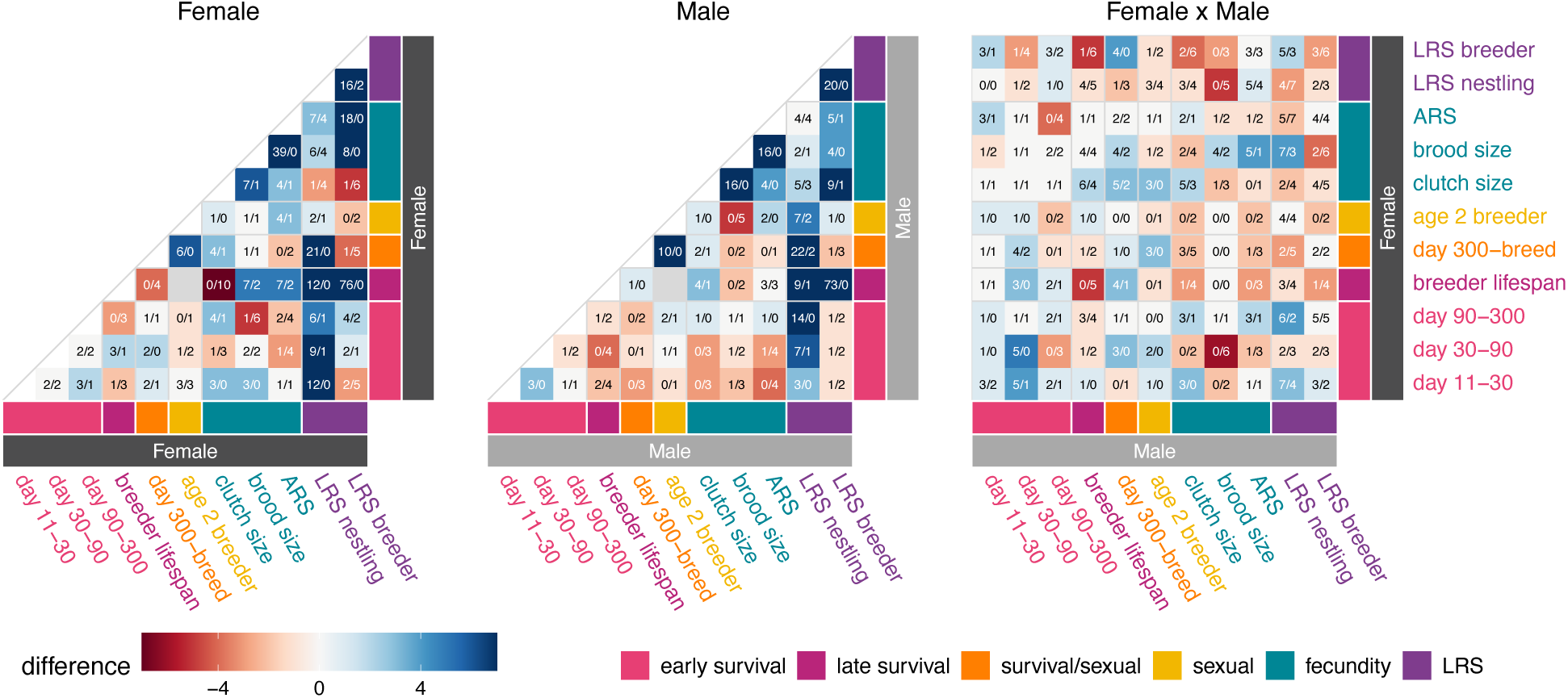
Concordant and antagonistic SNPs in each pair of analyses. Text in each box indicates concordant/antagonistic SNP counts. Cell color reflects the difference between these two counts, with the maximum absolute difference set to 7. We LD-pruned SNPs within each pairwise comparison (R^2^ < 0.2).

**Extended Data Figure 7.**
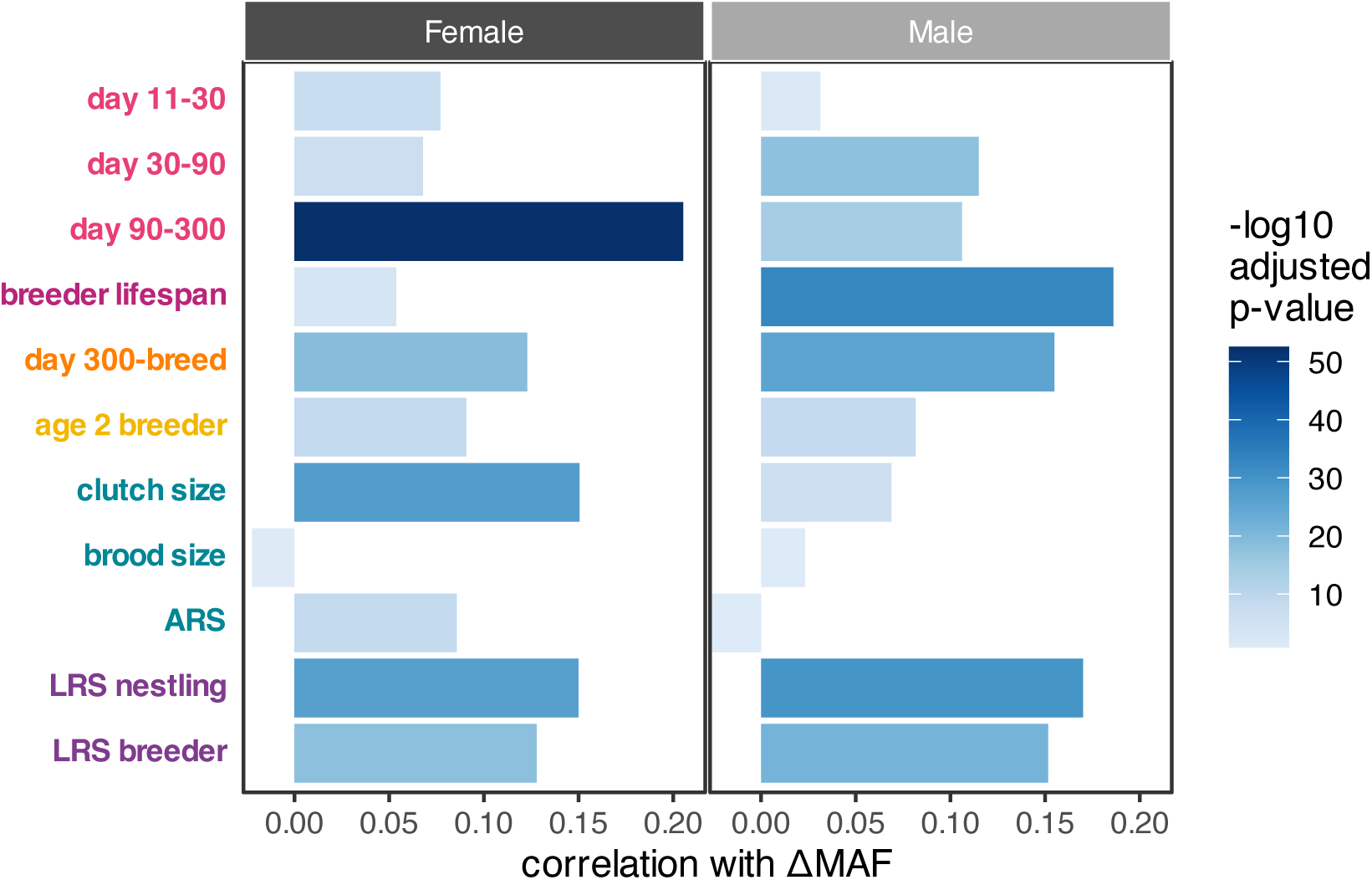
Correlation of effect size *z*-scores with allele frequency change (ΔMAF) estimates. Spearman correlation between effect size *z*-scores for each fitness measure analysis and ΔMAF estimates for 7,998 LD-pruned SNPs. Correlation *P*-values were generated via block jackknife (100 blocks). All Benjamini-Hochberg adjusted p-values are < 0.01 except the four smallest magnitude correlations: female brood size (adjusted *P* = 0.150), male day 11-30 (adjusted *P* = 0.036), male brood size (adjusted *P* = 0.098), and male ARS (adjusted *P* = 0.100).

**Extended Data Table 1:** Fitness measures used in this study with sample sizes and modeling information.

| Fitness component | Fitness measure | Description | Model family | Sample size |  |
| --- | --- | --- | --- | --- | --- |
|  |  |  |  | Female | Male |
| early survival | day 11-30 | survival from day 11 to day 30 | binomial | 1109 | 1082 |
|  | day 30-90 | survival from day 30 to day 90 | binomial | 663 | 673 |
|  | day 90-300 | survival from day 90 to day 300 | binomial | 497 | 520 |
| late survival | breeder lifespan | lifespan of breeders in years | Gaussian | 220 | 240 |
| survival/sexual | day 300-breed | survival from day 300 to becoming a breeder | binomial | 346 | 353 |
| sexual | age 2 breeder | breeder status at age 2 | binomial | 217 | 251 |
| fecundity | clutch size | number of eggs in first annual clutch | Gaussian | 755 | 770 |
|  | brood size | number of hatchlings in first annual clutch | Gaussian | 814 | 821 |
|  | ARS | total number of day 11 offspring produced annually | Gaussian | 808 | 822 |
| LRS | LRS nestling | total number of day 11 offspring produced in lifetime by locally born individuals | Poisson | 1148 | 1210 |
|  | LRS breeder | total number of day 11 offspring produced in lifetime by breeders (including immigrants) | Poisson | 220 | 240 |

**Extended Data Table 2:** SNPs significantly associated with a fitness component measure based on a per-analysis Benjamini-Hochberg adjusted *P*-value < 0.1.

| Sex | Component | Fitness measure | Chr | Position | MAF | Effect size | p-value | Adjusted p-value | Genotype model | Annotation | Gene name | LD group |
| --- | --- | --- | --- | --- | --- | --- | --- | --- | --- | --- | --- | --- |
| Female | Early Survival | day 11-30 | 11 | 1135002 | 0.14 | -0.69 | 6e-06 | 0.04 | Additive | 5'UTR | GAS8 |  |
| Female | Early Survival | day 11-30 | 14 | 5110333 | 0.22 | 0.62 | 4e-06 | 0.04 | Additive | intergenic_region |  |  |
| Female | Fecundity | clutch size | 12 | 20348267 | 0.39 | -0.42 | 2e-05 | 0.06 | Major dom | synonymous_variant | BHLHE40 |  |
| Female | Fecundity | clutch size | 14 | 8152838 | 0.21 | -0.74 | 1e-05 | 0.06 | Major dom | intergenic_region |  | 2 |
| Female | Fecundity | clutch size | 14 | 8308102 | 0.30 | -0.45 | 1e-05 | 0.06 | Major dom | intergenic_region |  | 2 |
| Female | Fecundity | brood size | 2 | 16592340 | 0.09 | 0.74 | 1e-05 | 0.06 | Minor dom | intergenic_region |  |  |
| Female | Fecundity | brood size | 2 | 166251531 | 0.16 | -1.54 | 8e-06 | 0.04 | Major dom | intergenic_region |  |  |
| Female | Fecundity | brood size | 26 | 3488936 | 0.14 | 0.64 | 3e-06 | 0.03 | Minor dom | intron_variant | ETV7 |  |
| Female | Fecundity | brood size | 9 | 13895723 | 0.25 | -1.04 | 1e-06 | 0.01 | Major dom | intergenic_region |  |  |
| Female | Fecundity | ARS | 10 | 11490217 | 0.48 | -0.62 | 2e-05 | 0.07 | Major dom | intergenic_region |  |  |
| Female | Fecundity | ARS | 13 | 1317138 | 0.33 | 0.81 | 3e-05 | 0.07 | Major dom | intron_variant | AFF4 |  |
| Female | Fecundity | ARS | 15 | 8677516 | 0.45 | -0.65 | 2e-05 | 0.07 | Major dom | intergenic_region |  |  |
| Female | Fecundity | ARS | 5 | 13634212 | 0.45 | -0.63 | 1e-05 | 0.07 | Major dom | downstream_gene_variant | PAK6 | 1 |
| Female | Fecundity | ARS | 5 | 13637967 | 0.45 | 0.59 | 7e-06 | 0.08 | Minor dom | downstream_gene_variant | PLCB2 | 1 |
| Male | Late Survival | breeder lifespan | 3 | 7078262 | 0.38 | 2.81 | 9e-07 | 0.01 | Major dom | intergenic_region |  |  |
| Male | Fecundity | ARS | 26 | 1451604 | 0.38 | -0.54 | 7e-06 | 0.08 | Minor dom | upstream_gene_variant | MYOG |  |

Abbreviations: Chr: chromosome; MAF: minor allele frequency in indicated analysis; Major dom: major allele dominance; Minor dom: minor allele dominance LD group: SNPs in LD ( $R^2 > 0.2$ ) are indicated by numeric grouping

**Extended Data Table 3:** SNPs significantly associated with a measure of LRS based on a per-analysis Benjamini-Hochberg adjusted *P*-value < 0.1.

| Sex | Fitness measure | Chr | Position | MAF | Effect size | p-value | Adjusted p-value | Genotype model | Annotation | Gene name | LD group |
| --- | --- | --- | --- | --- | --- | --- | --- | --- | --- | --- | --- |
| Female | LRS nestling | 17 | 7492863 | 0.23 | 2.08 | 5e-06 | 0.05 | Major dom | intron_variant | Vav2 |  |
| Male | LRS nestling | 11 | 18554667 | 0.19 | 2.81 | 8e-07 | 0.01 | Major dom | upstream_gene_variant | CDH11 |  |
| Male | LRS nestling | 2 | 168699161 | 0.15 | 2.90 | 1e-05 | 0.06 | Major dom | intron_variant | NRBP2 |  |
| Male | LRS nestling | 3 | 20575803 | 0.27 | 1.59 | 3e-05 | 0.10 | Major dom | intron_variant | FSHR |  |
| Male | LRS breeder | 19 | 9841527 | 0.28 | -0.66 | 9e-06 | 0.09 | Minor dom | downstream_gene_variant | TADA2A |  |
| Male | LRS breeder | 4 | 78933950 | 0.26 | 1.18 | 7e-06 | 0.04 | Major dom | intron_variant | RAB11FIP5 |  |

Abbreviations: Chr: chromosome; MAF: minor allele frequency in indicated analysis; Major dom: major allele dominance; Minor dom: minor allele dominance LD group: SNPs in LD ( $R^2 > 0.2$ ) are indicated by numeric grouping

## Notes

### Competing Interest Statement

The authors have declared no competing interest.

## References

1 Johnson, T. & Barton, N. Theoretical models of selection and mutation on quantitative traits. Philosophical Transactions of the Royal Society B: Biological Sciences 360, 1411–1425 (2005). 10.1098/rstb.2005.1667

2 Charlesworth, B. Causes of natural variation in fitness: Evidence from studies of Drosophila populations. Proceedings of the National Academy of Sciences 112, 1662–1669 (2015). 10.1073/pnas.1423275112

3 Lewontin, R. C. The Genetic Basis of Evolutionary Change. (Columbia University Press, 1974).

4 Zajitschek, F. & Connallon, T. Antagonistic pleiotropy in species with separate sexes, and the maintenance of genetic variation in life-history traits and fitness. Evolution 72, 1306–1316 (2018). 10.1111/evo.13493

5 Woolfenden, G. E. & Fitzpatrick, J. W. The Florida Scrub Jay - Demography of a cooperative-breeding bird. (Princeton University Press, 1984).

6 Dobzhansky, T. A review of some fundamental concepts and problems of population genetics. Cold Spring Harbor Symposia on Quantitative Biology 20, 1–15 (1955). 10.1101/sqb.1955.020.01.003

7 Charlesworth, B. & Hughes, K. A. in Evolutionary genetics: from molecules to morphology (eds R. S. Singh & C. B. Krimbas) 369–391 (Cambridge University Press, 2000).

8 Orr, H. A. Fitness and its role in evolutionary genetics. Nature Reviews Genetics 10, 531–539 (2009). 10.1038/nrg2603

9 Bonnet, T. et al. Genetic variance in fitness indicates rapid contemporary adaptive evolution in wild animals. Science 376, 1012–1016 (2022). 10.1126/science.abk0853

10 Ruzicka, F. et al. A century of theories of balancing selection. bioRxiv, 2025.2002.2012.637871 (2025). 10.1101/2025.02.12.637871

11 Curtsinger, J. W., Service, P. M. & Prout, T. Antagonistic Pleiotropy, Reversal of Dominance, and Genetic Polymorphism. The American Naturalist 144, 210–228 (1994). 10.1086/285671

12 Rose, M. R. Life history evolution with antagonistic pleiotropy and overlapping generations. Theoretical Population Biology 28, 342–358 (1985). 10.1016/0040-5809(85)90034-6

13 Maklakov, A. A. & Chapman, T. Evolution of ageing as a tangle of trade-offs: energy versus function. Proceedings of the Royal Society B: Biological Sciences 286, 20191604 (2019). 10.1098/rspb.2019.1604

14 Brown, K. E. & Kelly, J. K. Antagonistic pleiotropy can maintain fitness variation in annual plants. Journal of Evolutionary Biology 31, 46–56 (2018). 10.1111/jeb.13192

15 Carter, A. J. R. & Nguyen, A. Q. Antagonistic pleiotropy as a widespread mechanism for the maintenance of polymorphic disease alleles. BMC Medical Genetics 12, 160 (2011). 10.1186/1471-2350-12-160

16 Charmantier, A., Perrins, C., McCleery, R. H. & Sheldon, B. C. Quantitative genetics of age at reproduction in wild swans: Support for antagonistic pleiotropy models of senescence. Proceedings of the National Academy of Sciences 103, 6587–6592 (2006). 10.1073/pnas.0511123103

17 Flatt, T. Survival costs of reproduction in Drosophila. Experimental Gerontology 46, 369–375 (2011). 10.1016/j.exger.2010.10.008

18 Morton, A. J., Skillings, E. A., Wood, N. I. & Zheng, Z. Antagonistic pleiotropy in mice carrying a CAG repeat expansion in the range causing Huntington’s disease. Scientific Reports 9, 37 (2019). 10.1038/s41598-018-37102-8

19 Johnston, S. E. et al. Life history trade-offs at a single locus maintain sexually selected genetic variation. Nature 502, 93–95 (2013). 10.1038/nature12489

20 Foerster, K. et al. Sexually antagonistic genetic variation for fitness in red deer. Nature 447, 1107–1110 (2007). 10.1038/nature05912

21 Barson, N. J. et al. Sex-dependent dominance at a single locus maintains variation in age at maturity in salmon. Nature 528, 405–408 (2015). 10.1038/nature16062

22 Mérot, C., Llaurens, V., Normandeau, E., Bernatchez, L. & Wellenreuther, M. Balancing selection via life-history trade-offs maintains an inversion polymorphism in a seaweed fly. Nature Communications 11, 670 (2020). 10.1038/s41467-020-14479-7

23 Prout, T. in Evolutionary genetics: from molecules to morphology (eds R. S. Singh & C. B. Krimbas) 157–181 (Cambridge University Press, 2000).

24 Kidwell, J. F., Clegg, M. T., Stewart, F. M. & Prout, T. Regions of stable equilibria for models of differential selection in the two sexes under random mating. Genetics 85, 171–183 (1977). 10.1093/genetics/85.1.171

25 Connallon, T. & Chenoweth, S. F. Dominance reversals and the maintenance of genetic variation for fitness. PLOS Biology 17, e3000118 (2019). 10.1371/journal.pbio.3000118

26 Tellier, A., Villaréal, L. M. M. A. & Giraud, T. Antagonistic pleiotropy may help population-level selection in maintaining genetic polymorphism for transmission rate in a model phytopathogenic fungus. Heredity 98, 45–52 (2007). 10.1038/sj.hdy.6800902

27 Wong, H. W. S. & Holman, L. Pleiotropic fitness effects across sexes and ages in the Drosophila genome and transcriptome. Evolution 77, 2642–2655 (2023). 10.1093/evolut/qpad163

28 Mumme, R. L., Bowman, R., Pruett, M. S. & Fitzpatrick, J. W. Natal territory size, group size, and body mass affect lifetime fitness in the cooperatively breeding Florida Scrub-Jay. The Auk 132, 634–646 (2015). doi:10.1642/AUK-14-258.1

29 Aguillon, S. M. et al. Deconstructing isolation-by-distance: The genomic consequences of limited dispersal. PLOS Genetics 13, e1006911 (2017). 10.1371/journal.pgen.1006911

30 Efron, B. The Jackknife, the Bootstrap and Other Resampling Plans. (Society for Industrial and Applied Mathematics, 1982).

31 Chen, N., Cosgrove, E. J., Bowman, R., Fitzpatrick, J. W. & Clark, A. G. Genomic Consequences of Population Decline in the Endangered Florida Scrub-Jay. Current Biology 26, 2974–2979 (2016). 10.1016/j.cub.2016.08.062

32 Fitzpatrick, J. W. & Bowman, R. in Cooperative Breeding in Vertebrates (eds W. D. Koenig & J. L. Dickinson) 77-96 (Cambridge University Press, 2016).

33 Summers, J. et al. Context-Dependent Fitness Outcomes of Helping in the Cooperatively-Breeding Florida Scrub-Jay Aphelocoma coerulescens. bioRxiv (2025). 10.1101/2025.07.21.666021

34 Henderson, C. R. Applications of linear models in animal breeding. (University of Guelph, 1984).

35 Kruuk, L. B. E. Estimating genetic parameters in natural populations using the ‘animal model’. Philosophical Transactions of the Royal Society of London. Series B: Biological Sciences 359, 873–890 (2004). 10.1098/rstb.2003.1437

36 Falconer, D. S. & Mackay, T. F. C. Introduction to Quantitative Genetics. 4th edn, (Longman, 1996).

37 Williams, T. D. Mechanisms Underlying the Costs of Egg Production. BioScience 55, 39–48 (2005). 10.1641/0006-3568(2005)055[0039:MUTCOE]2.0.CO;2

38 Kruuk, L. E. et al. Heritability of fitness in a wild mammal population. Proc Natl Acad Sci U S A 97, 698–703 (2000). 10.1073/pnas.97.2.698

39 Merilä, J. & Sheldon, B. C. Genetic architecture of fitness and nonfitness traits: empirical patterns and development of ideas. Heredity 83, 103–109 (1999). 10.1046/j.1365-2540.1999.00585.x

40 Price, T. & Schluter, D. On the low heritability of life-history traits. Evolution 45, 853–861 (1991). 10.1111/j.1558-5646.1991.tb04354.x

41 Beavis, W. D. in Proceedings of the Forty-Ninth Annual Corn & Sorghum Industry Research Conference. (ed American Seed Trade Association) 250–266 (American Seed Trade Association).

42 Beavis, W. D. in Molecular Dissection of Complex Traits (ed A. H. Paterson) (CRC Press, 1998).

43 Crouch, D. J. M. & Bodmer, W. F. Polygenic inheritance, GWAS, polygenic risk scores, and the search for functional variants. Proc Natl Acad Sci U S A 117, 18924–18933 (2020). 10.1073/pnas.2005634117

44 Duntsch, L. et al. Polygenic basis for adaptive morphological variation in a threatened Aotearoa | New Zealand bird, the hihi (Notiomystis cincta). Proceedings of the Royal Society B: Biological Sciences 287, 20200948 (2020). 10.1098/rspb.2020.0948

45 Santure, A. W. et al. Replicated analysis of the genetic architecture of quantitative traits in two wild great tit populations. Molecular Ecology 24, 6148–6162 (2015). 10.1111/mec.13452

46 Bérénos, C. et al. Heterogeneity of genetic architecture of body size traits in a free-living population. Molecular Ecology 24, 1810–1830 (2015). 10.1111/mec.13146

47 Gauzere, J. et al. A polygenic basis for birth weight in a wild population of red deer (Cervus elaphus). G3 Genes|Genomes|Genetics 13, jkad018 (2023). 10.1093/g3journal/jkad018

48 Boyle, E. A., Li, Y. I. & Pritchard, J. K. An Expanded View of Complex Traits: From Polygenic to Omnigenic. Cell 169, 1177–1186 (2017). 10.1016/j.cell.2017.05.038

49 Di, C. & Lohmueller, K. E. Revisiting Dominance in Population Genetics. Genome Biology and Evolution 16, evae147 (2024). 10.1093/gbe/evae147

50 Woolfenden, G. E. & Fitzpatrick, J. W. Dominance in the Florida Scrub Jay. The Condor: Ornithological Applications 79, 1–12 (1977). 10.2307/1367524

51 Pettay, J. E., Kruuk, L. E. B., Jokela, J. & Lummaa, V. Heritability and genetic constraints of life-history trait evolution in preindustrial humans. Proceedings of the National Academy of Sciences 102, 2838–2843 (2005). 10.1073/pnas.0406709102

52 Whitlock, M. C. & Agrawal, A. F. Purging the genome with sexual selection: reducing mutation load through selection on males. Evolution 63, 569–582 (2009). 10.1111/j.1558-5646.2008.00558.x

53 Sultanova, Z., García-Roa, R. & Carazo, P. Condition-dependent mortality exacerbates male (but not female) reproductive senescence and the potential for sexual conflict. Journal of Evolutionary Biology 33, 1086–1096 (2020). 10.1111/jeb.13636

54 Bonduriansky, R. & Chenoweth, S. F. Intralocus sexual conflict. Trends in Ecology & Evolution 24, 280–288 (2009). 10.1016/j.tree.2008.12.005

55 Woolfenden, G. E. & Fitzpatrick, J. W. in Ecological aspects of social evolution: birds and mammals (eds D. Rubenstein & R.W. Wrangham) 97-107 (Princeton University Press, 1986).

56 Walsh, B. & Blows, M. W. Abundant Genetic Variation + Strong Selection = Multivariate Genetic Constraints: A Geometric View of Adaptation. Annual Review of Ecology, Evolution, and Systematics 40, 41–59 (2009).

57 Etterson, J. R. & Shaw, R. G. Constraint to Adaptive Evolution in Response to Global Warming. Science 294, 151–154 (2001). 10.1126/science.1063656

58 Connallon, T. & Clark, A. G. Antagonistic versus nonantagonistic models of balancing selection: characterizing the relative timescales and hitchhiking effects of partial selective sweeps. Evolution 67, 908–917 (2013). 10.1111/j.1558-5646.2012.01800.x

59 Santure, A. W. & Garant, D. Wild GWAS—association mapping in natural populations. Molecular Ecology Resources 18, 729–738 (2018). 10.1111/1755-0998.12901

60 Wolf, J. B., Brodie Iii, E. D., Cheverud, J. M., Moore, A. J. & Wade, M. J. Evolutionary consequences of indirect genetic effects. Trends in Ecology & Evolution 13, 64–69 (1998). 10.1016/S0169-5347(97)01233-0

61 Bijma, P. The quantitative genetics of indirect genetic effects: a selective review of modelling issues. Heredity 112, 61–69 (2014). 10.1038/hdy.2013.15

62 Kardos, M. & Luikart, G. The Genetic Architecture of Fitness Drives Population Viability during Rapid Environmental Change. The American Naturalist 197, 511–525 (2021). 10.1086/713469

63 Nguyen, T. N. et al. Whole-genome sequencing across space and time reveals impact of population decline and reduced gene flow in Florida Scrub-Jays. bioRxiv, 2024.2011.2005.622154 (2024). 10.1101/2024.11.05.622154

64 Chen, N. et al. Allele frequency dynamics in a pedigreed natural population. Proceedings of the National Academy of Sciences 116, 2158–2164 (2019). 10.1073/pnas.1813852116

65 Shah, S. S. et al. Lifetime fitness benefits of breeding site fidelity and low costs of inbreeding permit inbreeding tolerance in an avian cooperative breeder. bioRxiv, 2024.2009.2021.614049 (2024). 10.1101/2024.09.21.614049

66 Suh, Y. H., Pesendorfer, M. B., Tringali, A., Bowman, R. & Fitzpatrick, J. W. Investigating social and environmental predictors of natal dispersal in a cooperative breeding bird. Behavioral Ecology 31, 692–701 (2020). 10.1093/beheco/araa007

67 Quinn, J. S., Woolfenden, G. E., Fitzpatrick, J. W. & White, B. N. Multi-locus DNA fingerprinting supports genetic monogamy in Florida Scrub-Jays. Behavioral Ecology and Sociobiology 45, 1–10 (1999).

68 Townsend, A. K., Bowman, R., Fitzpatrick, J. W., Dent, M. & Lovette, I. J. Genetic monogamy across variable demographic landscapes in cooperatively breeding Florida Scrub-Jays. Behavioral Ecology 22, 464–470 (2011). 10.1093/beheco/arq227

69 Purcell, S. et al. PLINK: A Tool Set for Whole-Genome Association and Population-Based Linkage Analyses. The American Journal of Human Genetics 81, 559–575 (2007). 10.1086/519795

70 Driscoll, R. M. H. et al. Allele frequency dynamics under sex-biased demography and sex-specific inheritance in a pedigreed jay population. Genetics 227, iyae075 (2024). 10.1093/genetics/iyae075

71 Romero, F. G. et al. A new high-quality genome assembly and annotation for the threatened Florida Scrub-Jay (Aphelocoma coerulescens). G3 Genes|Genomes|Genetics 14, jkae232 (2024). 10.1093/g3journal/jkae232

72 Cingolani, P. et al. A program for annotating and predicting the effects of single nucleotide polymorphisms, SnpEff. Fly 6, 80–92 (2012). 10.4161/fly.19695

73 Chen, H. et al. Control for Population Structure and Relatedness for Binary Traits in Genetic Association Studies via Logistic Mixed Models. The American Journal of Human Genetics 98, 653–666 (2016). 10.1016/j.ajhg.2016.02.012

74 Amadeu, R. R., Garcia, A. A. F., Munoz, P. R. & Ferrão, L. F. V. AGHmatrix: genetic relationship matrices in R. Bioinformatics 39, btad445 (2023). 10.1093/bioinformatics/btad445

75 Schafer, J., et al. corpcor: Efficient Estimation of Covariance and (Partial) Correlation. (2021).

76 Hadfield, J. D. MCMC Methods for Multi-Response Generalized Linear Mixed Models: The MCMCglmm R Package. Journal of Statistical Software 33, 1–22 (2010). 10.18637/jss.v033.i02

77 de Villemereuil, P. Quantitative genetic methods depending on the nature of the phenotypic trait. Annals of the New York Academy of Sciences 1422, 29–47 (2018). 10.1111/nyas.13571

78 R: A Language and Environment for Statistical Computing (R Foundation for Statistical Computing, 2021).

