## Supplemental Text for "Genome-wide associations of fitness components reveal antagonistic pleiotropy and sexual conflict in the Florida Scrub-Jay"

### Supplementary Text

*Extra-pair paternity (EPP)*

Previous studies have established that Florida Scrub-Jays are almost strictly monogamous[1,2], but our large-scale genotyping allowed a more thorough check for extra-pair paternity (EPP)[3]. We used PLINK[4] and PedCheck[5] to test for Mendelian inconsistencies and found low rates of EPP[3]. Of the nests from mapped territories in 1989-1992 and 2003-2015 for which we had genotype data for both breeders and at least one nestling, we found evidence of EPP at 31 out of 576 nests (5.4%). We excluded nests with EPP from our analyses.

*Environmental covariates*

We considered many potential fixed effects in our models to control for variation in characteristics of the cohort, habitat, individual, natal nest environment, climate, and fire history. A full list of variables considered and their descriptions can be found in Supplemental Table 1.

For cohort effects, we considered annual variation in phenology, breeding opportunities, and population density. We used the first quartile of the distribution of annual hatch dates for the first clutch (in Julian days) to estimate population-wide changes in the timing of breeding each year ("annual tract-wide hatch date"). All territories in the study population are mapped each year, allowing us to count the number of territories with a breeding vacancy each year and calculate density as the inverse of the mean territory size for a given year. We also considered the number of male or female adults as a measure of population size changes over time.

Using fine-scale maps of habitat composition and annual territory maps, we calculated the proportion of each territory composed of roads or scrub habitat in ArcGIS[6]. To estimate food abundance over time, we used historical data on acorn counts, an important food source for Florida Scrub-Jays when invertebrates are less abundant in the winter[7]. The number of acorns for multiple marked plants across the study tract are counted each fall[8], allowing us to calculate the total acorn count across all five oak species present in our study site each year.

Individuals in our population were sexed based on behavioral field observations prior to the collection of blood for sexing using standard molecular methods[9] and sex assignments were confirmed with genomic data. We calculated genomic-based inbreeding coefficients for each individual using the $F^{III}$ estimator in PLINK v1.9 (option -ibc)[4]. Immigrants are defined as individuals who were not born in our study area.

Given documented preference for short dispersal distances[10,11], we considered the number of breeding vacancies for each sex within 1-3 territories of the natal territory. We also considered the presence or number of helpers at the nest as well as the age, inbreeding coefficient, and immigrant status of the parents. We know that breeding experience has detectable effects on reproductive success in Florida Scrub-Jays[12,13], so we considered the breeding experience of individual parents and of the pair. We estimated pairwise relatedness of parents using the proportion of the genome shared identical-by-descent between breeding pairs calculated in PLINK v1.9 (option -genome).

We obtained Southern Oscillation Index (SOI) data from the Australian Government Bureau of Meteorology (ftp://ftp.bom.gov.au/anon/home/ncc/www/sco/soi/soiplaintext.html). Archbold Biological Station maintains a weather station on site that collects information on daily rainfall (in inches), drought index (a metric of soil moisture levels), and temperature (https://www.archbold-station.org/gis-and-data-mgmt/). We summarized low temperatures as both the mean minimum temperature and the sum of degrees (Fahrenheit) below freezing during a focal time interval. We considered SOI, rainfall, drought index, and temperature measures for multiple time intervals relevant to the specific fitness measure.

Finally, as Florida Scrub-Jay life history is tightly linked to the natural fire cycle[14], we also considered different measures of fire history for each territory. Archbold Biological Station performs prescribed burning to help maintain the scrub habitat and keeps fire history maps at a fine spatial resolution. We used ArcGIS to calculate the proportion of the territory that was burned in the previous year or in the last 1-9 years as well as the mean number of years since fire for each territory each year.

*Phenotypic correlations between fitness components*

We looked at phenotypic correlations between our various measures of late survival, breeder status, fecundity, and LRS (Supplemental Figure 1). We note that many pairwise comparisons involving early survival measures are not possible because individuals must survive to breed in order to reproduce. We also excluded within-sex comparisons between breeder status at age 2 and breeder lifespan, as any individual with a breeder lifespan of only 2 years had attained breeder status by age 2, forcing a negative correlation between these two measures. While most fitness measures are positively correlated, as expected, we found that breeder status and brood size were significantly negatively correlated in females (Spearman's rho = -0.23, $P = 0.02$). In general, observed positive correlations of genome-wide $z$-scores of effect sizes between LRS and different fitness components (Figure 4) mirrored phenotypic correlations (Supplemental Figure 1).

*Fixed effects associated with fitness measures*

We applied a subsampling approach[15] to select significant environmental covariates for each fitness measure in females and males separately (Extended Data Figure 1, Supplemental Figure 2). As our focus here was on selecting potential confounding variables for downstream association analyses and not on estimating the effects of each variable on fitness, we do not expect our results to perfectly match results of previous studies investigating inbreeding depression or the effects of territory size or number of helpers on fitness [3,12-14,16]. However, the variables selected in our analyses were largely consistent with our understanding of Florida Scrub-Jay biology.

We found evidence of inbreeding depression for multiple fitness measures. High inbreeding was associated with lower female day11-30 and day90-300 survival as well as lower male $LRS_{nestling}$. Previous studies of inbreeding depression in this population found significant associations of inbreeding with lower day90-300 survival, male and female breeder lifespan, female LRS, and hatching success[3]. Notably, we found that relatedness of the breeding pair was positively associated with clutch size, which could suggest that more inbred pairs are compensating for lower hatching success by increasing clutch size.

Our variable selection results also recovered known fitness benefits of large territories[12,13]. Breeding territory size was positively associated with ARS for both sexes. Natal territory size was positively associated with breeder status at age 2 for males, consistent with previous results showing that large natal territory size increased the probability a male became a breeder[12]. We found that the number of helpers at the nest was negatively associated with ARS and clutch size in both sexes, potentially indicating a cost of having helpers due to increased competition for resources within a given territory.

*Excluding morphometrics and incubation date*

Our fitness measure effect size estimates include any potential contributions from variation in individual morphometric measures and incubation date. Though we know that size and incubation date are associated with variation in survival and reproductive success in Florida Scrub-Jays, respectively[12,13], we did not include these measures as potential covariates because we wanted our effect size estimates to reflect all potential genetic contributions to variation in fitness, including effects of alleles that affect fitness indirectly by influencing size or nesting behavior. We quantified the additive genetic variance for three individual morphometric measures (nestling weight, juvenile weight, and juvenile tarsus length) and incubation date (in Julian dates) using the same approach used for the fitness measures. Briefly, we performed variable selection and then fitted animal models using MCMCglmm[17]. We found that weight and tarsus traits in both sexes were affected by brood size, experience of the breeders, and other attributes of the natal territory and climate. Environmental covariates explained 37% and 41% of the observed variation in incubation date in females and males, respectively; selected variables included whether a pair has bred together previously and the proportion of the territory that consists of roads (Supplemental Figure 3). We observed non-zero additive genetic variance in all three morphometric variables in both sexes except male nestling weight (Supplemental Figure 4), indicating that these traits are heritable. While we did not find significant additive genetic variance for incubation date, we still excluded it from our potential covariates as timing of reproduction is known to have a genetic component in multiple birds[18-20] and a large proportion of variance in this trait could be explained by other covariates (Supplemental Figure 4). Both the annual tract-wide hatch date and whether the breeding pair is a new pair are strong predictors of incubation date and were considered as covariates in our fecundity models.

*Note on modeling approaches*

For LRS, we used different modeling approaches for estimating additive genetic variance and for genome-wide association analysis. We assumed that LRS follows a zero-inflated Poisson (ZIP) distribution (Figure 1C) and followed the approach of Bonnet et al.[21] to fit Bayesian animal models using MCMCglmm for heritability estimation. These models also allowed direct estimation of uncertainty. Unfortunately, MCMCglmm is not a computationally feasible option for fitting the thousands of models required for genome-wide association analysis. Furthermore, ZIP regression models have separate coefficient estimates for the zero-inflated and Poisson portions, rather than a single coefficient estimate, making comparisons to other effect size estimates difficult. Thus, we used GMMAT[22] with a Poisson family to perform genome-wide association analysis for LRS.

In our genome-wide association analysis, we considered three genotype models: additive (*AA* = 0, *Aa* = 1, *aa* = 2), major allele dominance (*AA* = *Aa* = 0, *aa* = 1), and minor allele dominance (*AA* = 0, *Aa* = *aa* = 1). We did not attempt to explicitly quantify dominance effects here, as detecting nonadditive effects requires much larger sample sizes[23]. Instead, our goal was to maximally capture the effects of a given allele, and the dominance encodings employed herein have greater potential of capturing an effect with both additive and dominance components than an additive model alone.

*The effect of gene flow*

Previous work found high but decreasing levels of gene flow into our study population over time, with incoming immigrants exhibiting lower levels of heterozygosity compared to locally born individuals[3]. To account for any potential fitness differences between immigrants and locally born individuals, we included immigrant status of individuals or parents as a potential covariate in all relevant analyses. Whether or not an individual was an immigrant was a significant predictor of female $LRS_{breeder}$, suggesting there are fitness differences between female residents and immigrants in our study population. Immigrant status was not a significant predictor of overall male fitness or any specific fitness component.

Dispersal in Florida Scrub-Jays is female-biased[10,11], leading to a marked difference in the number of female vs male immigrants: there were 84 female vs. 25 male immigrants in the fecundity data sets, and 58 female vs. 14 male immigrants in the $LRS_{breeder}$ and breeder lifespan data sets. We repeated our analyses of genome-wide effect sizes excluding immigrant observations and obtained very similar results (Supplemental Figure 5). Thus, we do not believe the different trends observed for females and males in Figure 4 are driven by sex-biases in immigration rate.

*Signatures of antagonistic pleiotropy without environmental covariates*

To ensure that the patterns we found were not driven by different fixed effects included in each sex-specific fitness measure model, we repeated our analysis of polygenic signals of pleiotropy using effect size *z*-scores obtained from genome-wide association models that only included the GRM and other biologically-relevant random effects (as shown in Figure 2). We note that in some cases, the variance explained by fixed effects is likely absorbed by one or more random effects when we remove fixed effects from the models. Genome-wide correlations of *z*-scores obtained from models without any fixed effects (Supplemental Figure 6) showed similar patterns of within-sex trade-offs between fitness components as the original analysis (Figure 4). For the between-sex comparison, removing fixed effects revealed more antagonism between $LRS_{breeder}$ in one sex and fecundity measures in the other sex, as well as sexual antagonism between $LRS_{nestling}$. Overall, the inclusion of different environmental covariates in different fitness measure models does not drive the observed prevalence of genetic trade-offs within and between sexes.

*Signatures of antagonistic pleiotropy among candidate SNPs*

We repeated our analysis of polygenic signals of pleiotropy using only LD-pruned SNPs with $P < 0.01$ for at least one of the considered fitness measures (84-190 SNPs per comparison, mean 147). Correlations of *z*-scores for LD-pruned candidate SNPs (Supplemental Figure 7) are largely consistent with patterns found in genome-wide correlations using all LD-pruned SNPs (Figure 4), just with fewer significant comparisons. We still found evidence of genome-wide trade-offs between early- and late-survival (specifically between survival from day 90 to day 300 and breeder survival in females and between survival from day 11 to day 30 and survival from day 300 to breed in males) as well as between survival and fecundity both within and between sexes. In males, effect sizes for survival from day 30 to day 90 are negatively correlated with those for all three fecundity measures (Supplemental Figure 7). We did not find any evidence of sexual conflict when comparing effect sizes for the same fitness component between sexes but found multiple cases of negative correlations between female survival and male sexual selection and fecundity.

### References


1 Quinn, J. S., Woolfenden, G. E., Fitzpatrick, J. W. & White, B. N. Multi-locus DNA fingerprinting supports genetic monogamy in Florida Scrub-Jays. *Behavioral Ecology and Sociobiology* **45**, 1-10 (1999).

2 Townsend, A. K., Bowman, R., Fitzpatrick, J. W., Dent, M. & Lovette, I. J. Genetic monogamy across variable demographic landscapes in cooperatively breeding Florida Scrub-Jays. *Behavioral Ecology* **22**, 464-470 (2011). https://doi.org/10.1093/beheco/arq227

3 Chen, N., Cosgrove, E. J., Bowman, R., Fitzpatrick, J. W. & Clark, A. G. Genomic Consequences of Population Decline in the Endangered Florida Scrub-Jay. *Current Biology* **26**, 2974-2979 (2016). https://doi.org/10.1016/j.cub.2016.08.062

4 Purcell, S. *et al.* PLINK: A Tool Set for Whole-Genome Association and Population-Based Linkage Analyses. *The American Journal of Human Genetics* **81**, 559-575 (2007). https://doi.org/https://doi.org/10.1086/519795

5 O'Connell, J. R. & Weeks, D. E. PedCheck: A Program for Identification of Genotype Incompatibilities in Linkage Analysis. *The American Journal of Human Genetics* **63**, 259-266 (1998). https://doi.org/https://doi.org/10.1086/301904

6 ArcGIS Release 10 (Environmental Systems Research Institute, Redlands, CA, 2011).

7 DeGange, A. R., Fitzpatrick, J. W., Layne, J. N. & Woolfenden, G. E. Acorn Harvesting by Florida Scrub Jays. *Ecology* **70**, 348-356 (1989). https://doi.org/10.2307/1937539

8 Pesendorfer, M. B. *et al.* Fire history and weather interact to determine extent and synchrony of mast-seeding in rhizomatous scrub oaks of Florida. *Philosophical Transactions of the Royal Society B: Biological Sciences* **376** (2021). https://doi.org/10.1098/rstb.2020.0381
9 Fridolfsson, A.-K. & Ellegren, H. A Simple and Universal Method for Molecular Sexing of Non-Ratite Birds. *Journal of Avian Biology* **30**, 116-121 (1999). https://doi.org/10.2307/3677252
10 Aguillon, S. M. *et al.* Deconstructing isolation-by-distance: The genomic consequences of limited dispersal. *PLOS Genetics* **13**, e1006911 (2017). https://doi.org/10.1371/journal.pgen.1006911
11 Suh, Y. H., Pesendorfer, M. B., Tringali, A., Bowman, R. & Fitzpatrick, J. W. Investigating social and environmental predictors of natal dispersal in a cooperative breeding bird. *Behavioral Ecology* **31**, 692-701 (2020). https://doi.org/10.1093/beheco/araa007
12 Mumme, R. L., Bowman, R., Pruett, M. S. & Fitzpatrick, J. W. Natal territory size, group size, and body mass affect lifetime fitness in the cooperatively breeding Florida Scrub-Jay. *The Auk* **132**, 634-646 (2015). https://doi.org/10.1642/AUK-14-258.1
13 Woolfenden, G. E. & Fitzpatrick, J. W. *The Florida Scrub Jay - Demography of a cooperative-breeding bird*. (Princeton University Press, 1984).
14 Fitzpatrick, J. W. & Bowman, R. in *Cooperative Breeding in Vertebrates* (eds W. D. Koenig & J. L. Dickinson) 77-96 (Cambridge University Press, 2016).
15 Efron, B. *The Jackknife, the Bootstrap and Other Resampling Plans*. (Society for Industrial and Applied Mathematics, 1982).
16 Summers, J. *et al.* Context-Dependent Fitness Outcomes of Helping in the Cooperatively-Breeding Florida Scrub-Jay Aphelocoma coerulescens. *bioRxiv* (2025). https://doi.org/10.1101/2025.07.21.666021
17 Hadfield, J. D. MCMC Methods for Multi-Response Generalized Linear Mixed Models: The MCMCglmm R Package. *Journal of Statistical Software* **33**, 1 - 22 (2010). https://doi.org/10.18637/jss.v033.i02
18 Jones, C. V., Regan, C. E., Cole, E. F., Firth, J. A. & Sheldon, B. C. Shared environmental similarity between relatives influences heritability of reproductive timing in wild great tits. *Evolution* **79**, 220-231 (2025). https://doi.org/10.1093/evolut/qpae155
19 Teplitsky, C., Mills, J. A., Yarrall, J. W. & Merilä, J. Indirect genetic effects in a sex-limited trait: the case of breeding time in red-billed gulls. *Journal of Evolutionary Biology* **23**, 935-944 (2010). https://doi.org/10.1111/j.1420-9101.2010.01959.x
20 Sheldon, B. C., Kruuk, L. E. B. & Merila, J. Natural selection and inheritance of breeding time and clutch size in the collared flycatcher. *Evolution* **57**, 406-420 (2003). https://doi.org/https://doi.org/10.1111/j.0014-3820.2003.tb00274.x

21 Bonnet, T. *et al.* Genetic variance in fitness indicates rapid contemporary adaptive evolution in wild animals. *Science* **376**, 1012-1016 (2022). https://doi.org/10.1126/science.abk0853
22 Chen, H. *et al.* Control for Population Structure and Relatedness for Binary Traits in Genetic Association Studies via Logistic Mixed Models. *The American Journal of Human Genetics* **98**, 653-666 (2016). https://doi.org/10.1016/j.ajhg.2016.02.012
23 Palmer, D. S. *et al.* Analysis of genetic dominance in the UK Biobank. *Science* **379**, 1341-1348 (2023). https://doi.org/10.1126/science.abn8455

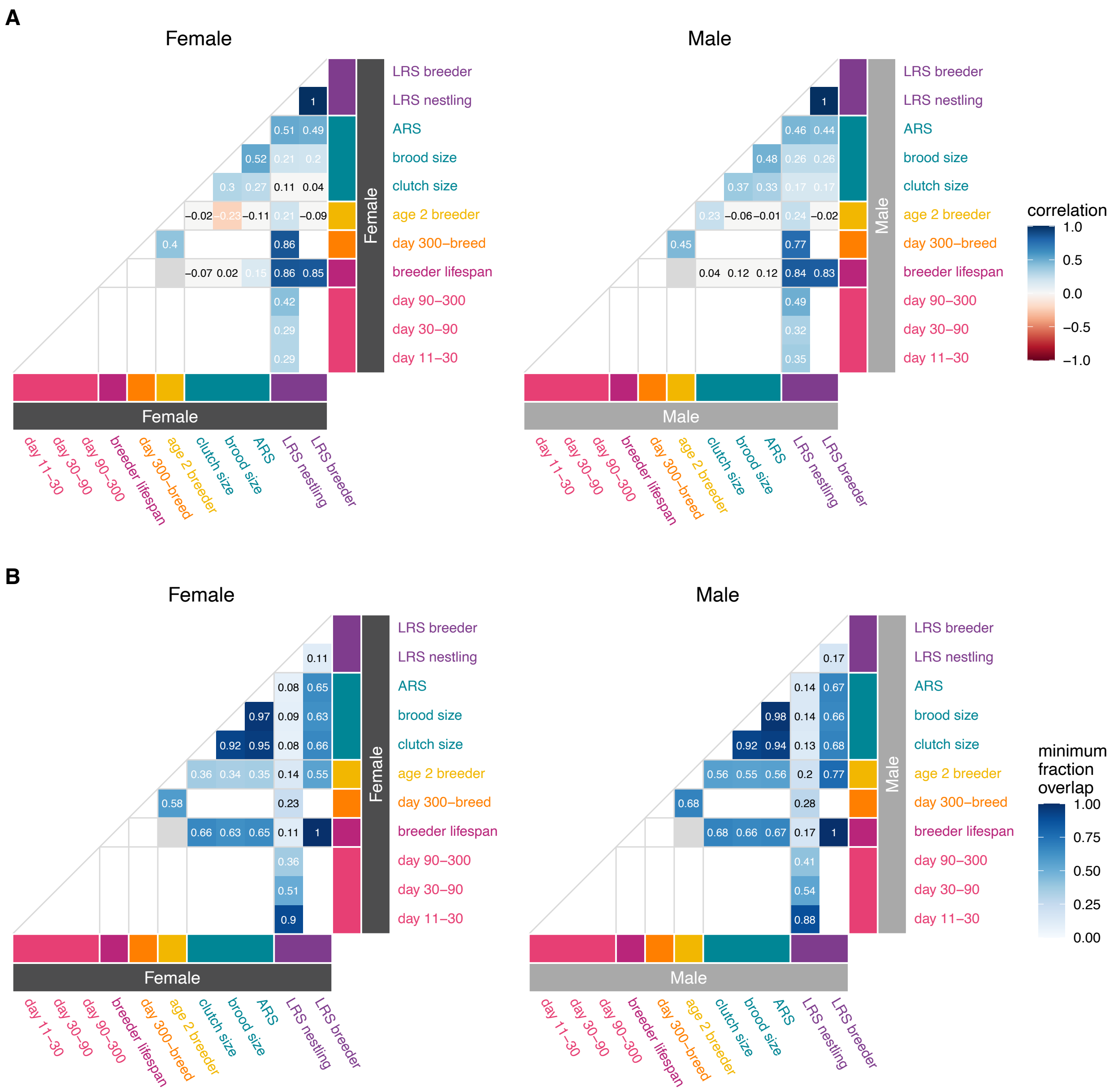


**Supplemental Figure 1. Phenotypic correlations and sample overlap across fitness measures. (A)** Spearman correlation between each pair of fitness measures. All correlations for valid comparisons are shown. Significant correlations (adjusted *P*-value < 0.1) are shown in white text in cells colored by the correlation coefficient, while non-significant correlations are shown in black text. Empty cells reflect invalid comparisons (cases where all samples with values for both measures had the same fitness value for one of the two measures). **(B)** Sample overlap between a given pair of fitness measures. For each pair of fitness measures, we computed the fraction of samples in one analysis included in the other analysis, and vice versa, and the minimum of the two values is plotted. Only values corresponding to non-empty comparisons in **(A)** are plotted.

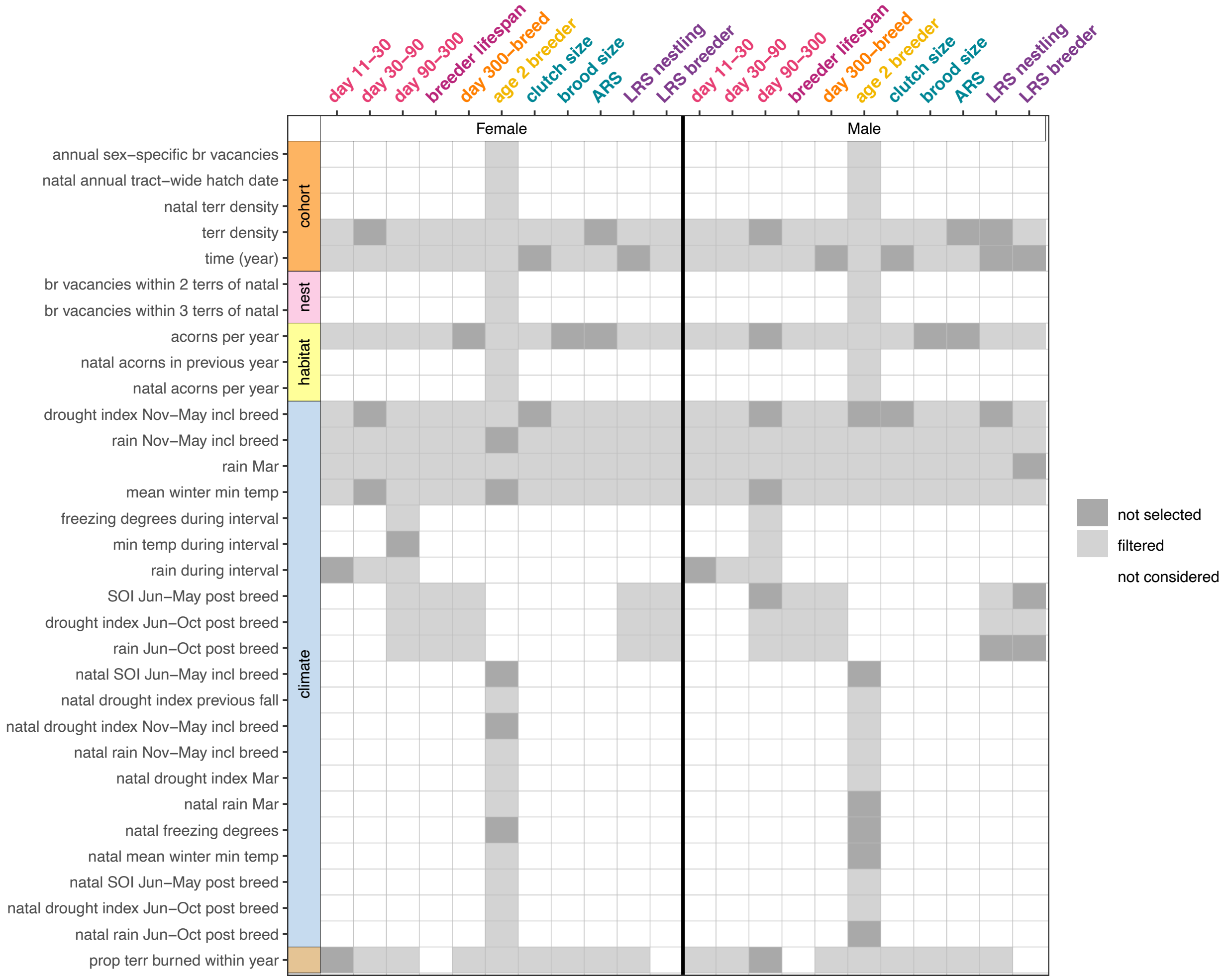


**Supplemental Figure 2. Additional fixed effects that were never selected.** These variables included characteristics of the cohort (orange), natal nest environment (pink), habitat (yellow), climate (blue), and fire history (brown). Gray shading indicates variables that were removed due to earlier filtering steps (light gray) or at the final variable selection step (dark gray).

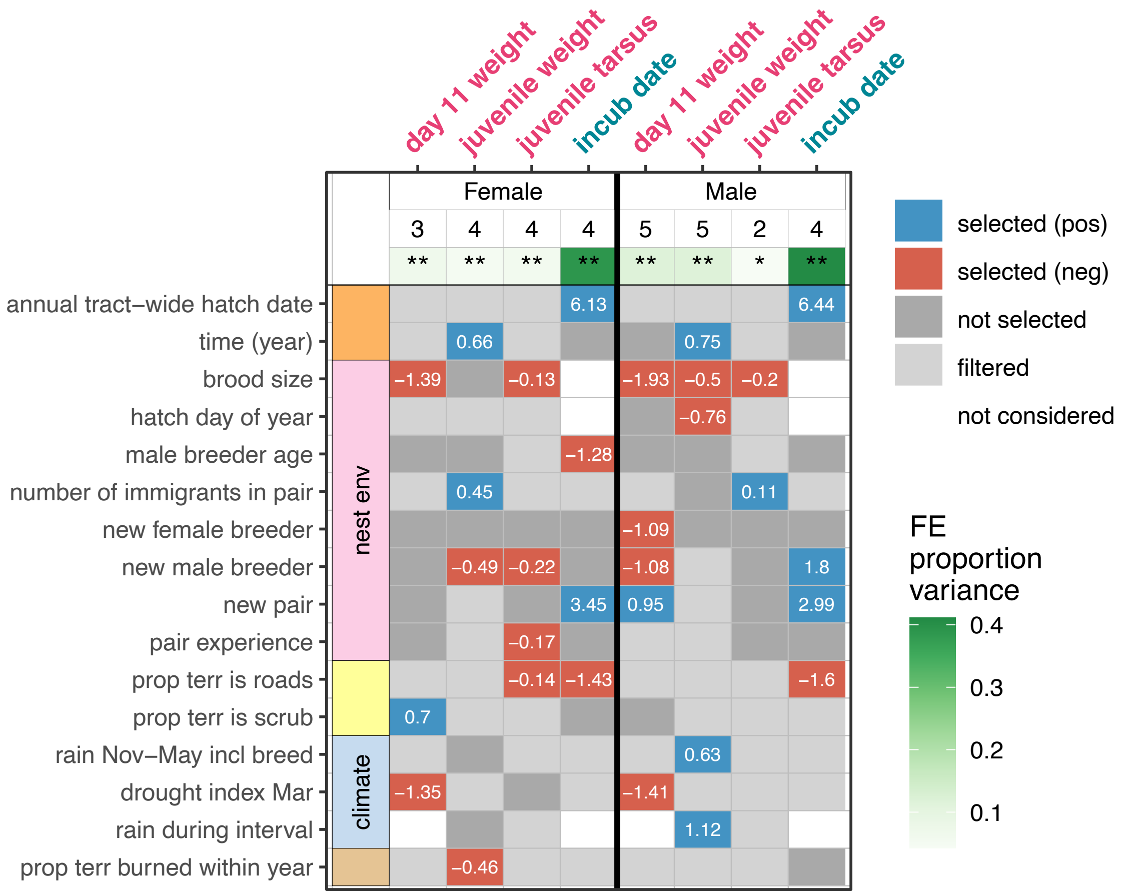


**Supplemental Figure 3. Fixed effects selected for analyses of morphometric traits and incubation date.** Numeric values indicate the regression coefficient in the final model for each fitness measure in females and males. We considered several different potential covariates (fixed effects; FE), including characteristics of the cohort (orange), individual (none selected here), natal nest environment (pink), habitat (yellow), climate (blue), and fire history (brown). Gray shading indicates variables that were removed due to earlier filtering steps (light gray) or at the final variable selection step (dark gray). The number of variables included in downstream analyses and the posterior mode proportion of phenotypic variance explained by these variables derived from Bayesian animal models (Supplemental Figure 4) are displayed across the top. Single and double asterisks indicate that the lower bound of the 95% credible interval for the corresponding variance estimate is greater than 0.001 or 0.01, respectively.

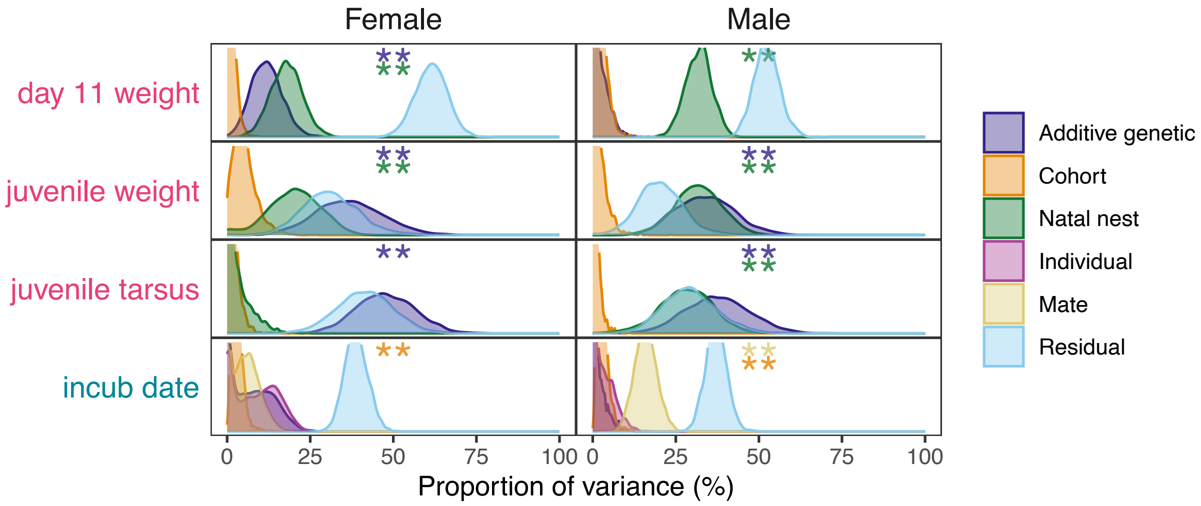


**Supplemental Figure 4. Components of variance in female and male morphometrics and incubation date.** Posterior distributions of additive genetic variance (purple) as well as variance due to cohort (orange) or natal nest (green) effects. Cohort effects refer to breeding cohort for incubation date and natal cohort for morphometric traits. For incubation date, individual and mate identity are shown in pink and yellow, respectively. Single and double asterisks indicate that the lower bound of the 95% credible interval of corresponding variance estimate is greater than 0.001 or 0.01, respectively.

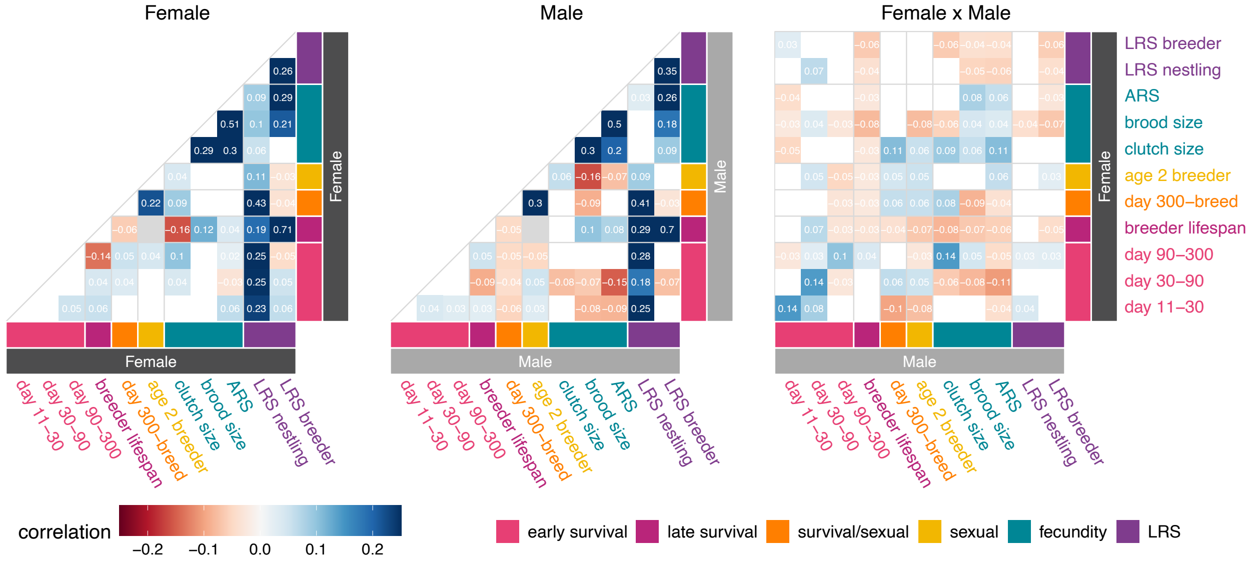


**Supplemental Figure 5. Polygenic signatures of pleiotropy excluding immigrants.** We excluded immigrants from all data sets and repeated the full analysis presented in Figure 4. Pairwise Spearman correlation between effect size *z*-scores for 7,998 LD-pruned SNPs for each pair of fitness measures within females (left), within males (center), or between sexes (right). Correlation *P*-values were generated via block jackknife (100 blocks). Only correlations with Benjamini-Hochberg adjusted *P* < 0.1 are shown. The correlation color scale ranges from -0.25 to 0.25.

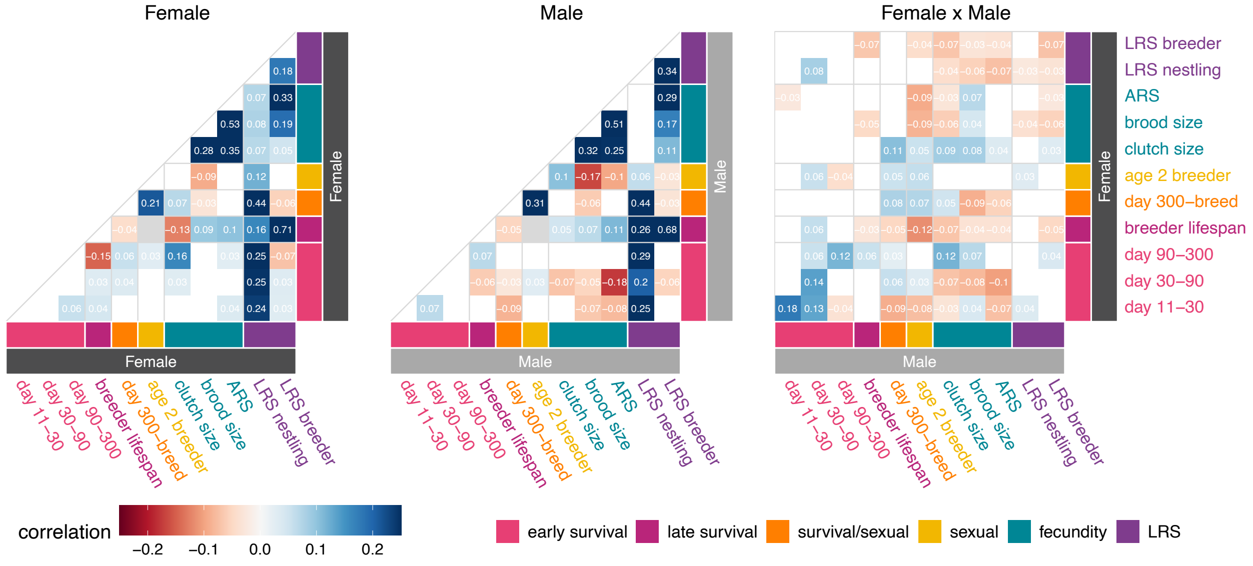


**Supplemental Figure 6. Polygenic signatures of pleiotropy without consideration of environmental covariates.** We repeated the full analysis presented in Figure 4 without fixed effects in the models. Pairwise Spearman correlation between effect size *z*-scores for 7,998 LD-pruned SNPs for each pair of fitness measures within females (left), within males (center), or between sexes (right). Correlation *P*-values were generated via block jackknife (100 blocks). Only correlations with Benjamini-Hochberg adjusted $P < 0.1$ are shown. The correlation color scale ranges from -0.25 to 0.25.

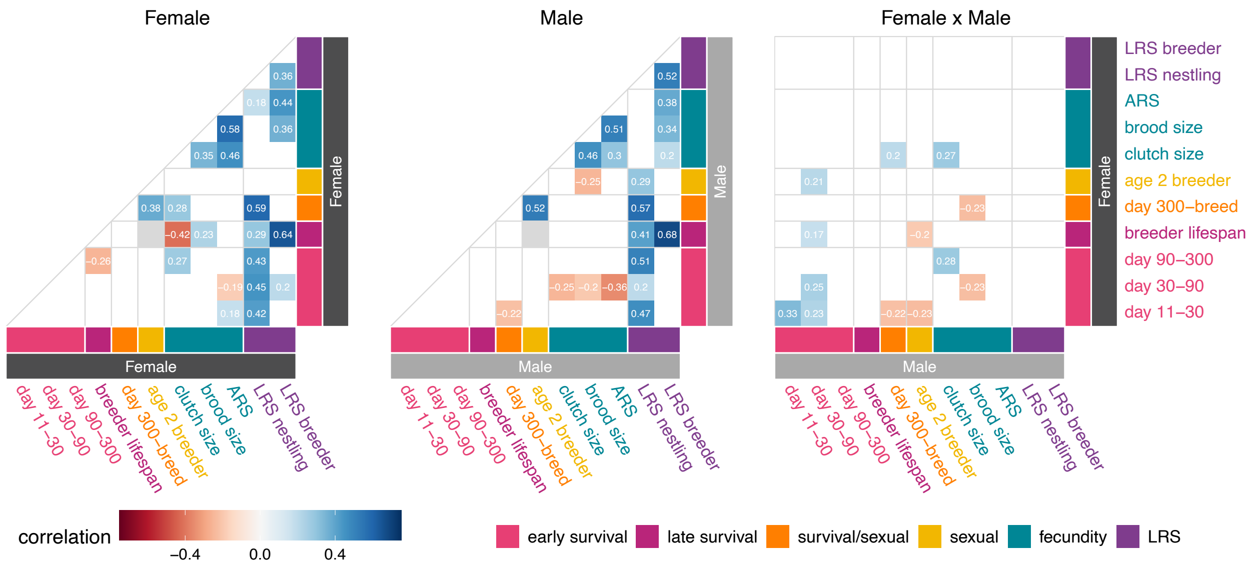


**Supplemental Figure 7. Polygenic signatures of pleiotropy limited to candidate SNPs.** Pairwise Spearman correlation coefficients between effect size estimates for LD-pruned candidate SNPs (*p* < 0.01 in either analysis) for each pair of fitness measures within females (left), within males (center), or between sexes (right). The number of SNPs included in each comparison ranged from 84-190 (median 147). Only correlations with Benjamini-Hochberg adjusted *P* < 0.1 are shown. The correlation color scale ranges from -0.75 to 0.75.

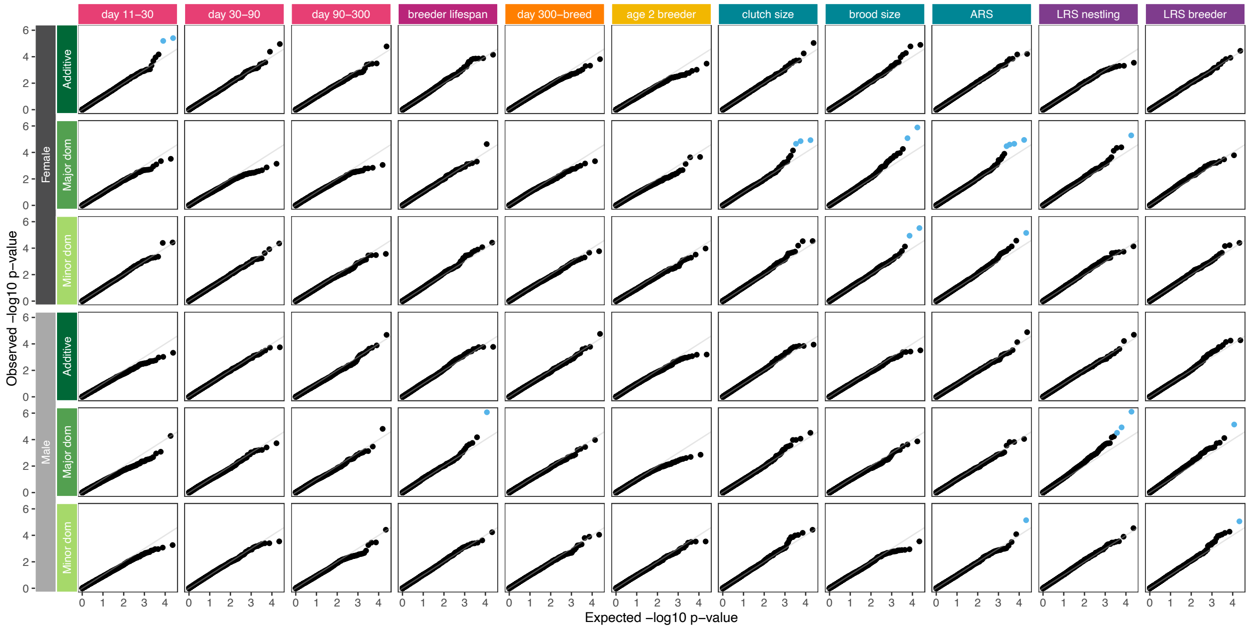


**Supplemental Figure 8. Quantile-quantile plots for all genome-wide association analyses.** Observed vs. expected -log10 *P*-values for each genome-wide association analysis, with significant SNPs (adjusted $P < 0.1$) shown in blue. Green labels indicate genotype model. Major dom: major allele dominance. Minor dom: minor allele dominance.

**Supplemental Table 1: Environmental variables considered in models of fitness components and total fitness.**

| Category | Variable | Description |
|---|---|---|
| Cohort | annual tract-wide hatch date | annual first quartile value of study tract-wide first clutch hatch day of year |
| | annual sex-specific br vacancies | count of annual sex-specific breeding vacancies |
| | sex-specific adult population size | annual sex-specific adult population size |
| | terr density | 1 / annual mean territory size |
| | time (year) | year as quantitative (numeric) value |
| | natal annual tract-wide hatch date | natal year annual tract-wide hatch date |
| | natal terr density | natal year terr density |
| Habitat | acorns in previous year | acorns per year of previous year |
| | acorns per year | annual sum of acorns per stem at posts surveyed 1988:2017 (measured study tract-wide annually, not per territory) |
| | prop terr is roads | proportion of territory area covered by roads |
| | prop terr is scrub | proportion of territory area covered by oak scrub |
| | terr size | territory size in hectares |
| | natal acorns in previous year | natal year acorns in previous year |
| | natal acorns per year | natal year acorns per year |
| Individual | inbreeding coef | inbreeding coefficient estimated from the SNP data using plink1.9 –ibc Fhat3 |
| | is immigrant | 1 if individual is an immigrant, 0 otherwise |
| | sex | molecularly determined sex of the individual |
| Nest environment | br vacancies within 1 terr of natal | count of sex-specific breeding vacancies (previous year breeder not observed after May 1 of current breeding season) with territory centroids within one territory distance (median study-wide sqrt(territory size)) of natal territory centroid |
| | br vacancies within 2 terrs of natal | count of sex-specific breeding vacancies with territory centroids within 2x territory distance of natal territory centroid |
| | br vacancies within 3 terrs of natal | count of sex-specific breeding vacancies with territory centroids within 3x territory distance of natal territory centroid |
| | any helpers | 1 if helper count $>$ 0, 0 otherwise |
| | helper count | count of non-breeder individuals age 1 or older observed at nest territory |
| | hatch day of year | hatch date in day of year (Julian date + 1) |
| | brood size | number of hatchlings in natal nest |
| | female breeder age | female breeder age in years, immigrants assumed age 2 at first breeding season observation |
| | female breeder inbreeding coef | inbreeding coef of the female breeder, estimated from the SNP data using plink1.9 –ibc Fhat3 |
| | female breeder is immigrant | 1 if female breeder in an immigrant, 0 otherwise |
| | male breeder age | male breeder age in years, immigrants assumed age 2 at first breeding season observation |
| | male breeder inbreeding coef | inbreeding coef of the male breeder, estimated from the SNP data using plink1.9 –ibc Fhat3 |
| | male breeder is immigrant | 1 if male breeder in an immigrant, 0 otherwise |
| | number of immigrants in pair | count of immigrants in breeding pair (0, 1, or 2) |
| | pair IBD | identity-by-descent of breeding pair, calculated by plink 1.9 –genome |
| | is new breeder | 1 if breeder is new breeder, 0 otherwise |
| | mate is new breeder | 1 if mate is new breeder, 0 otherwise |
| | new female breeder | 1 if female breeder is new, 0 otherwise |
| | new male breeder | 1 if male breeder is new, 0 otherwise |
| | new pair | 1 if pair is new, 0 otherwise |
| | pair experience | count of previous breeding seasons for a given pair |

### Supplemental Table 1: continued

| Category | Variable | Description |
|---|---|---|
| Climate | SOI Jun-May incl breed | mean Southern Oscillation Index June-May preceding and including breeding season |
| | drought index previous fall | mean drought index Sep-Nov preceding breeding season |
| | rain previous fall | sum rain (inches) Sep-Nov preceding breeding season |
| | drought index Nov-May incl breed | mean drought index Nov-May (preceding and including breeding season) |
| | rain Nov-May incl breed | sum rain (inches) Nov-May (preceding and including breeding season) |
| | drought index Mar | mean drought index March |
| | rain Mar | sum rain (inches) in March |
| | freezing degrees | sum of degrees (Fahrenheit) below freezing for daily minimum temperatures Nov-April preceding and overlapping breeding season |
| | mean winter min temp | mean minimum temperatures (degrees Fahrenheit) Dec-Feb in winter preceding breeding season |
| | drought index during interval | mean drought index during the fitness measure time interval |
| | freezing degrees during interval | sum of degrees (Fahrenheit) below freezing for daily minimum temperatures during the fitness measure interval |
| | max temp during interval | mean maximum temperatures (degrees Fahrenheit) during the fitness measure time interval |
| | min temp during interval | mean minimum temperatures (degrees Fahrenheit) during the fitness measure time interval |
| | rain during interval | sum rain (inches) during the fitness measure time interval |
| | SOI Jun-May post breed | mean Southern Oscillation Index June-May post breeding season |
| | drought index Jun-Oct post breed | mean drought index Jun-Oct post breed |
| | rain Jun-Oct post breed | sum rain (inches) Jun-Oct post breed |
| | natal SOI Jun-May incl breed | natal year SOI Jun-May incl breed |
| | natal drought index previous fall | natal year drought index previous fall |
| | natal rain previous fall | natal year rain previous fall |
| | natal drought index Nov-May incl breed | natal year drought index Nov-May incl breed |
| | natal rain Nov-May incl breed | natal year rain Nov-May incl breed |
| | natal drought index Mar | natal year drought index Mar |
| | natal rain Mar | natal year rain Mar |
| | natal freezing degrees | natal year freezing degrees |
| | natal mean winter min temp | natal year mean winter min temp |
| | natal SOI Jun-May post breed | natal year SOI Jun-May post breed |
| | natal drought index Jun-Oct post breed | natal year drought index Jun-Oct post breed |
| | natal rain Jun-Oct post breed | natal year rain Jun-Oct post breed |
| Fire | prop terr burned within year | proportion of territory burned in the year preceding a given observation |
| | prop terr burned in last 1 to 9 years | proportion of territory burned in the last one to nine years |
| | terr mean time since fire | mean years since fire over territory area |

In fecundity analyses, “Nest environment” variables refer to observed breeder/pair/nest;
for all other analyses, these refer to natal nest or parents.
In sexual selection analyses, “Fire” and “Habitat” variables refer to natal values.
